# Rapid Cortical Adaptation and the Role of Thalamic Synchrony During Wakefulness

**DOI:** 10.1101/2020.10.08.331660

**Authors:** Nathaniel C. Wright, Peter Y. Borden, Yi Juin Liew, Michael F. Bolus, William M. Stoy, Craig R. Forest, Garrett B. Stanley

## Abstract

Rapid sensory adaptation is observed across all sensory systems, and strongly shapes sensory percepts in complex sensory environments. Yet despite its ubiquity and likely necessity for survival, the mechanistic basis is poorly understood. A wide range of primarily in-vitro and anesthetized studies have demonstrated the emergence of adaptation at the level of primary sensory cortex, with only modest signatures in earlier stages of processing. The nature of rapid adaptation and how it shapes sensory representations during wakefulness, and thus the potential role in perceptual adaptation, is underexplored, as are the mechanisms that underlie this phenomenon. To address these knowledge gaps, we recorded spiking activity in primary somatosensory cortex (S1) and the upstream ventral posteromedial (VPm) thalamic nucleus in the vibrissa pathway of awake male and female mice, and quantified responses to whisker stimuli delivered in isolation and embedded in an adapting sensory background. We found that cortical sensory responses were indeed adapted by persistent sensory stimulation; putative excitatory neurons were profoundly adapted, and inhibitory neurons only modestly so. Further optogenetic manipulation experiments and network modeling suggest this largely reflects adaptive changes in synchronous thalamic firing combined with robust engagement of feedforward inhibition, with little contribution from synaptic depression. Taken together, these results suggest that cortical adaptation in the regime explored here results from changes in the timing of thalamic input, and the way in which this differentially impacts cortical excitation and feedforward inhibition, pointing to a prominent role of thalamic gating in rapid adaptation of primary sensory cortex.

**Significance Statement:** Rapid adaptation of sensory activity strongly shapes representations of sensory inputs across all sensory pathways over the timescale of seconds, and has profound effects on sensory perception. Despite its ubiquity and theoretical role in the efficient encoding of complex sensory environments, the mechanistic basis is poorly understood, particularly during wakefulness. In this study in the vibrissa pathway of awake mice, we show that cortical representations of sensory inputs are strongly shaped by rapid adaptation, and that this is mediated primarily by adaptive gating of the thalamic inputs to primary sensory cortex and the differential way in which these inputs engage cortical sub-populations of neurons.

## Introduction

Our experience of the world around us depends upon context. For instance, a noisy environment provides persistent sensory stimulation, which can adapt the representations and percepts of salient sensory features embedded within the environment. Rapid sensory adaptation describes such interactions between stimulus history and perception, spanning milliseconds to seconds (Whitmire and Stanley, 2016). A wealth of human psychophysical studies have documented perceptual adaptation in audition (Smith and Faulkner, 2006; Bestelmeyer et al., 2010; Erb et al., 2013), vision (Blakemore and Campbell, 1969; Blakemore and Nachmias, 1971; Anstis et al., 1998; Ghodrati et al., 2019), and somatosensation (Tannan et al., 2007), suggesting rapid sensory adaptation lends a vital flexibility to organisms tasked with surviving and thriving during rapid environmental changes.

Despite the ubiquity of rapid sensory adaptation across sensory pathways, and its likely relevance to function (Barlow, 1961), the neural basis has not been conclusively identified. A large body of (mostly in vitro and anesthetized) work has implicated the early sensory pathway; adaptation effects cascade toward and culminate in primary sensory cortex (Ganmor et al., 2010; Maravall et al., 2013; Lampl and Katz, 2017), and the nature of cortical adaptation is consistent with perceptual effects. In the vibrissa pathway of the anesthetized rodent for example, adaptation induced through whisker stimulation shapes the amplitude (Webber and Stanley, 2006, 2004; Boloori and Stanley, 2006; Maravall et al., 2007; Heiss et al., 2008; Boloori et al., 2010; Ganmor et al., 2010; Wang et al., 2010; Cohen-Kashi Malina et al., 2013; Ollerenshaw et al., 2014; Kheradpezhouh et al., 2017) and spatial extent (Ollerenshaw et al., 2014; Zheng et al., 2015) of responses to subsequent stimuli in primary somatosensory cortex (S1). Yet two important questions remain unanswered. First, what is the nature of cortical response adaptation during wakefulness, where baseline activity is elevated relative to the anesthetized state (Greenberg et al., 2008; Vizuete et al., 2012; Aasebø et al., 2017)? The thalamocortical pathway during wakefulness may be in a baseline “adapted” state that is relatively impervious to additional adaptation (Castro-Alamancos, 2004), and the neural basis for perceptual adaptation lies elsewhere, but this has not been rigorously tested. Second, if the circuit is subject to sensory adaptation during wakefulness, what are the underlying mechanisms? Previous anesthetized and in-vitro work implicates thalamocortical and/or intracortical synaptic depression (Castro-Alamancos and Oldford, 2002; Chung et al., 2002; Gabernet et al., 2005; Cruikshank et al., 2007, 2010; Cohen-Kashi Malina et al., 2013), which could explain why mean evoked rates are generally more adapted in cortex than in thalamus (Chung et al., 2002; Khatri et al., 2004; Gabernet et al., 2005). Yet adaptation also reduces thalamic synchrony (Wang et al., 2010; Ollerenshaw et al., 2014) and single-unit bursting (Whitmire et al., 2016) under anesthesia. Given the sensitivity of cortex to the timing of thalamic inputs (Usrey et al., 2000; Swadlow and Gusev, 2001; Bruno and Sakmann, 2006; Wang et al., 2010; Ollerenshaw et al., 2014), this suggests a potential role for thalamic spike timing in cortical adaptation.

Here, we address these unknowns by recording from and modeling S1 of the awake, head-fixed mouse during rapid sensory adaptation. We found that despite the relatively high level of baseline activity typical of wakefulness, putative excitatory neurons in S1 were profoundly adapted by background sensory stimulation. In particular, mean evoked firing rates, theoretical stimulus detectability, and synchronous cortical spiking were all significantly reduced in the adapted state, consistent with previously-observed changes in perceptual report (Ollerenshaw et al., 2014; Waiblinger et al., 2015). Several lines of evidence – including the recording of the thalamic inputs under a range of optogenetic controls and computational modeling – suggest this primarily reflected reduced synchronous thalamic firing and robust thalamically-driven feedforward inhibition in the adapted condition, with little contribution from thalamocortical and intracortical synaptic depression. Taken together, these results establish the role of the thalamocortical circuit in rapid adaptation during wakefulness, and implicate a more critical role of thalamic input than previously thought.

## Materials and Methods

All procedures were approved by the Institutional Animal Care and Use Committee at the Georgia Institute of Technology (Protocol Numbers A100223 and A100225), and were in agreement with guidelines established by the National Institutes of Health.

### Surgery

Experiments were carried out using C57BL/6J, Ai32xNR133 (Bolus et al., 2020) (Ai32 crossed with the NR133 Cre-recombinase driver line (Gerfen et al., 2013)), and Ai32xPV-Cre mice of both sexes. All mouse lines were purchased from Jackson Laboratories. Mice were induced with 5% isoflurane in an induction chamber, then transferred to a heating pad on a stereotaxic instrument, and maintained at 1 - 2% isoflurane for the remainder of the surgical procedure. A custom stainless steel headplate was fixed to the exposed skull with Metabond dental cement (Parkell, Inc.), exposed bone and tissue were then sealed with Metabond and either super-glue (Loctite 404; Henkel) or Vetbond Tissue Adhesive (3M). Metabond was used to fashion a well surrounding the left hemisphere. The well was filled with Kwik-Cast (World Precision Instruments, Inc.) and covered with a thin layer of Metabond, and the mouse was returned to its home cage. Mice were given pre- (buprenorphine) and post-operative (ketoprofen) analgesic, and were allowed to recover for three days before additional handling.

### Habituation

Three days after headplate implantation, mice were handled for at least 15 minutes, and then returned to their home cage. On subsequent days, mice were gradually habituated to head-fixation on a custom platform consisting of a tunnel with headpost clamps at one end. The first three daily habituation sessions lasted 15, 30, and 45 minutes respectively, but mice were returned to their home cage if they displayed signs of distress. We then gradually extended session durations until mice would tolerate at least 1.5 hours of fixation and whisker stimulation without signs of distress. Mice that did not meet these criteria were removed from the study, or used for anesthetized recordings (see below).

### Awake electrophysiological recordings

We recorded from five Ai32xNR133, four C57BL/6J (wild-type) and one Ai32xPV-cre awake mice of both sexes (up to three awake sessions per mouse). We used intrinsic optical signal imaging acquired under anesthesia to identify at least one putative principal column in S1. On the morning of the first recording session for each animal, we anesthetized the mouse as described above, and opened an approximately 500 micron × 500 micron craniotomy centered over a putative cortical column. When acquiring simultaneous VPm and S1 recordings, we opened a second craniotomy of similar size over the stereotactic coordinates for VPm (1.8 mm lateral from midline by 1.8 mm caudal from bregma) and slowly inserted either a single-channel tungsten electrode (2 MOhm, FHC) or 32-channel silicon probe (NeuroNexus A1×32-Poly3-5mm-25s-177) to a depth of approximately 3 mm. We adjusted the electrode depth while presenting continuous 10 Hz “sawtooth” stimulus trains to individual whiskers until we could identify a putative principal barreloid (by observing broad-waveform units with robust, short-latency, minimally-adapting sensory responses to stimulation of a single whisker, and at most comparatively weak responses to stimulation of surrounding whiskers (Brecht and Sakmann, 2002)), before slowly retracting the electrode/probe. We then covered exposed brain tissue with agarose, filled the well with Kwik-Cast, and allowed the mouse to recover in its home cage for at least two hours. After recovery, we head-fixed the awake mouse, removed the Kwik-Cast, filled the well with either saline, mineral oil, or agarose, and inserted an electrode/probe into each open craniotomy using a Luigs and Neumann manipulator. For S1 recordings, we inserted a multi-channel silicon probe (NeuroNexus) oriented 35 degrees from vertical. We used either a 32-channel linear (A1×32-5mm-25-177-A32), 32-channel “Poly3” (A1×32-Poly3-5mm-25s-177), or 64-channel, four-shank “Poly2” (A4×16-Poly2-5mm-23s-200-177) configuration probe. For VPm recordings, we inserted either a tungsten optoelectrode (2 Megaohm, FHC, with attached 200 micron optic fiber, Thorlabs), 32-channel silicon probe (A1×32-Poly3-5mm-25s-177) or 32-channel silicon optoelectrode (A1×32-Poly3-5mm-25s-177-OA32LP, with attached 105 micron optic fiber coupled to a 200 micron optic fiber, Thorlabs). Optic fibers were coupled to a 470 nm LED (M470F3, Thorlabs). When the barreloid we functionally identified during the anesthetized VPm mapping session was not topographically-aligned with the targeted S1 column, we referenced the (Coronal) Allen Brain Atlas to adjust the position of the VPm probe before descending. We inserted the probe(s) slowly to avoid excessive tissue dimpling, and waited at least 30 minutes after probe insertion to begin recording, to allow the tissue to settle. Continuous signals were acquired using a either a Cerebus (Blackrock Microsystems) or Tucker Davis Technologies acquisition system. Signals were amplified, filtered between 0.3 Hz and 7.5 kHz, and digitized at either 30 kHz or 24414.0625 Hz.

After the first recording session, we removed the probe(s), covered exposed tissue with agarose, and sealed the well with Kwik-Cast and a thin layer of Metabond. We obtained either two or three recording sessions (one per day) from each mouse using the original craniotomy, but each time targeting a different cortical column and barreloid.

### Anesthetized extracellular recordings

We recorded from three C57BL/6J (wild-type), four Ai32xPV-Cre, and three Ai32xNR133 mice of both sexes under isoflurane anesthesia. Mice were anesthetized and implanted with headplates, and we opened a single craniotomy (either approximately 500 microns × 500 microns, or 1 mm × 1 mm) over S1, as described above. In some cases, the principal column was first identified using intrinsic optical signal imaging, as described above. In other cases, we inserted a single tungsten electrode into the stereotactic coordinates for the center of S1, and defined the putative principal whisker to be that which evoked the largest LFP response. We then inserted a 4 × 16 silicon probe array (A4×16-Poly2-5mm-23s-200-177) to a depth of 700 microns. We oriented the probe to avoid blood vessels on the cortical surface. For a subset of these experiments, we obtained additional sessions by repeating the stimulation protocol using the whisker that evoked the maximum LFP response on a shank different from the first. For each such session, we determined the putative principal column off-line using white-noise-evoked spiking. For each shank, we summed single- and multi-unit (see below) spiking across all trials for the 1 s window preceding feature onset. We divided the across-trial mean white-noise-evoked response by the across-trial standard deviation of spontaneous spiking, and the shank with the largest resulting value was determined to correspond to the principal column.

### Anesthetized intracellular recordings

We recorded ongoing, sensory-evoked, and light-evoked subthreshold activity from four sensory- and light-responsive neurons in two mice using an Autopatcher system (Kodandaramaiah et al., 2016), as described in detail previously (Stoy et al., 2020). Briefly, we head-plated and identified the putative C2 column using intrinsic optical signal imaging in two isoflurane-anesthetized Ai32xNR133 transgenic mice, and opened a 1 mm × 1 mm craniotomy over the column, as described above. We then used an Autopatcher 1500 (Neuromatic Devices) to provide pressure and measure pipette resistance, and an algorithm based on these measurements to navigate around blood vessels in an automated fashion while the pipette descended through cortical tissue. Finally, we applied a recently-developed automated motion-compensation procedure (Stoy et al., 2020) for synchronizing the motion of the pipette tip to that of the targeted neuron prior to forming a seal. These experiments utilized Multiclamp 700B amplifiers (Molecular Devices), and signals were digitized at 20 kHz (cDAQ-9174, National Instruments), and were recorded in PClamp 10 in current-clamp mode.

### Whisker stimulation

We used a precise, computer-controlled galvanometer (Cambridge Technologies) with attached tube to stimulate individual whiskers (Whitmire et al., 2016; Waiblinger et al., 2018; Sederberg et al., 2019; Liew et al., 2020). The galvanometer was controlled using either a custom Matlab GUI and Simulink Real-Time (MathWorks), or the Real-time eXperiment Interface application (http://rtxi.org/), sampling at 1 kHz. We inserted the whisker into the tube, which was positioned approximately 10 mm from the whisker pad. We delivered “sawtooth” stimulus features (exponential rise and decay waveforms lasting approximately 17 ms, with reported velocity defined by the average over the 8.5 ms rising phase (Wang et al., 2010)) either in isolation, or embedded in frozen sensory white noise (i.e., white noise waveforms that were identical across trials). To generate the white noise waveforms, the value at each time-step was drawn from a Gaussian distribution (with standard deviation of one degree), and the resulting signal was lowpass-filtered at 100 Hz (3rd-order Butterworth (Waiblinger et al., 2015)). The white noise waveform around the feature waveform was dampened with an inverted Gaussian, with standard deviation 25.5 ms.

The stimulus conditions were randomized across trials. The stimulus consisted of 1.5 s of either white noise (“adapted” trials) or no white noise (“control” trials), with the onset of the embedded feature at 1 s. The inter-trial interval was a random value (drawn from a uniform distribution) between 2 and 3 s. In a subset of experiments, optogenetic stimulation of either VPm or thalamocortical terminals was randomly interleaved (see below). We typically obtained at least 100 trials per stimulus condition.

### Optogenetic stimulation

In a subset of acute anesthetized experiments in Ai32xNR133 transgenic mice, we stimulated thalamocortical terminals in S1 using blue (470 nm) light from an LED (ThorLabs), and recorded either spiking or subthreshold S1 responses. We positioned either a 200 or 400 micron optic fiber (ThorLabs) just above the exposed cortical surface, adjacent to the probe or patch pipette. Light pulses were either 10 ms or 15 ms in duration, and were delivered either in isolation or embedded in sensory white noise delivered to the whisker by the galvanometer. We titrated the light level at the beginning of each recording session to evoke cortical responses that were comparable in amplitude to those evoked by punctate whisker stimulation.

In a subset of awake experiments in Ai32xNR133 transgenic mice, we presented the above sensory stimulus protocol, in addition to a set of “LED” trials in which we delivered a step input of 470 nm light to VPm beginning 1 s before and ending 0.5 s after the delivery of an isolated sensory feature. The light was delivered via LED-coupled fiber attached to the electrode/probe (described above). We titrated the light level at the beginning of each recording session such that steady-state light-evoked firing rates in VPm (based on threshold crossings of high-pass-filtered voltage recordings) approximately matched those evoked by the white noise whisker stimulus.

### Videography and whisker motion analysis

In five awake recording sessions, we also recorded whisker videography using a CCD camera (DMK 21BU04.H USB 2.0 monochrome industrial camera, The Imaging Source, LLC). The face was illuminated using IR LEDs (860 nm, DigiKey), and the face opposite the galvanometer was imaged at 30 Hz. We used external triggers to acquire frames, and recorded these triggers in Synapse to synchronize frames to all other recorded signals. We analyzed the videos using the python implementation of FaceMap (www.github.com/MouseLand/FaceMap) (Stringer et al., 2019). For each video, we used the FaceMap GUI to select two regions of interest (ROIs): one that captured most of the whiskers, and a second of the nose. We then processed the videos for each ROI, which involved first calculating the “motion energy” of each frame (i.e., the absolute difference between the current and previous frame), and then computing the singular vectors of the motion energy. For each video, we confirmed that the time series of the resulting first SVDs for each ROI (which we call the “motion timeseries”) were qualitatively consistent with nose/whisker dynamics. We then calculated the relative whisker and nose motion as a function of the sensory stimulus for each recording session. To do this, we first calculated the across-time mean value of the motion timeseries in the 1 s preceding sensory feature onset for each trial, which we call the “mean motion”. We compared the mean motion values for the two stimulus conditions (“spontaneous” and “white noise”) using the Wilcoxon signed-rank test. For visualization, we then calculated the across-trial mean absolute motion value for the “spontaneous” and “white noise” conditions, and normalized to the “spontaneous” value.

### Spike-sorting

We sorted spikes off-line using KiloSort2 (https://github.com/MouseLand/Kilosort2) for clustering, and phy (https://github.com/cortex-lab/phy) for manual curation of clusters. During manual curation, clusters were either merged or separated based primarily on waveform distributions across the probe and cross-correlogram structure. We discarded as “noise” those clusters with across-instance mean waveform that did not resemble a characteristic spike on any channel. All remaining clusters were labeled as either single- or multi-units by downstream analysis (see below).

### Units retained for analysis

We labeled each curated cluster as either a single- or multi-unit based on the signal-to-noise ratio (SNR) and inter-spike interval (ISI) distribution. For each cluster and recorded spike, we calculated the absolute voltage difference between the trough and subsequent peak (VTP) on each channel. We defined the SNR to be the across-trial mean VTP divided by the across-trial standard deviation, for the channel on which the mean VTP was greatest. Additionally, we calculated “ISI violation percentage” for each cluster using the autocorrelogram (ACG). We defined the violation percentage to be the percentage of spikes in the 0 - 1 ms ACG bin. We then defined a well-isolated single-unit to be a cluster with SNR greater than 4.0, VTP greater than 50 mV, and ISI violation percentage below 1%. All other clusters were classified as multi-units. In our anesthetized recording sessions, we used an alternate probe configuration, which was somewhat less well-suited to obtaining well-isolated units. We therefore slightly relaxed our inter-spike interval violation constraints (to 1.5%) for defining “well-isolated” units, to yield more RS and FS cells from these datasets (see Methods). This did not qualitatively change our results. For S1, we further segregated single-units into regular- and fast-spiking units based on the mean waveform. Again using the channel on which the waveform was largest, we calculated the time from trough to subsequent peak (TTP). We classified S1 units as either fast-spiking (FS, putative inhibitory) or regular-spiking (RS, putative excitatory) neurons based on waveform width (McCormick et al., 1985; Niell and Stryker, 2010; Guo et al., 2017; Isett et al., 2018; Speed et al., 2019; Yu et al., 2019). Specifically, S1 units with TTP < 0.4 ms were classified as FS, and all others as RS. Waveforms were in general narrower for VPm units than for cortical units, consistent with previous work (Barthó et al., 2014). We therefore classified VPm “RS” cells as those with TTP > 0.3 ms, and excluded units with narrower waveforms, which likely originated from either the cell bodies or axon terminals of neurons in reticular thalamus (Barthó et al., 2014). For putative VPm units, we further required that the absolute peak of the PSTH of responses to isolated punctate stimuli occur between 2 and 10 ms of stimulus onset. Finally, when analyzing activity of single- and/or multi-units, we only included those units with at least 0.25 mean post-feature spikes per trial, and a significant change (p < 0.05, Wilcoxon signed-rank test) in firing rate after stimulus onset on control (unadapted) trials, using the weakest sawtooth stimulus delivered during that recording session and 50 ms (30 ms) pre- and post-stimulus windows for S1 (VPm).

### Experimental Design and Statistical Analyses

When comparing two sets of values (that were matched samples) across stimulus conditions, we used the Wilcoxon signed-rank test (implemented in Python using the wilcoxon function in the Scipy library), and Bonferroni-corrected the resulting p-values for multiple comparisons where applicable. When comparing two independent samples (e.g., normalized response rates for RS and FS cells), we used the Kruskal-Wallis test (implemented in Python using the kruskal function in the Scipy library).

For any analysis resulting in a single value for a given recording session calculated using all trials (e.g., mean synchronous spikes), we tested for significance of change across conditions by re-sampling trials with replacement, re-calculating the final value for the re-sampled pseudo-data, and calculating the 95% confidence intervals (Bonferroni-corrected if necessary) of the resulting distribution of values.

The number of cells and animals used to calculate each reported result is included in the text of the Results section and/or figure captions.

### Analyses

All analyses were performed using custom scripts in Python 3.0. The details of each analysis are presented below.

### ROC analysis

We calculated the theoretical detectability of sensory features for each significantly-responsive RS unit (see above) by applying ideal observer analysis (Wang et al., 2010; Ollerenshaw et al., 2014) to the “population response” distributions for baseline and feature-evoked activity. For this analysis, we excluded putative inhibitory neurons (FS units), as we were interested in interpreting the loss of excitatory drive from cortical neurons, likely to play a role in downstream signaling and thus relevant for percepts. For each unit, we calculated the firing rate in a 50 ms baseline (pre-feature) and post-feature window on each trial. For baseline activity, we used a window beginning 500 ms preceding feature onset, as the white noise was largely dampened in the 50 ms window immediately preceding the feature on adapted trials. We then calculated the across-trial mean (µ) and standard deviation (σ) of the observed firing rates, and generated “population” firing rate distributions, or samples drawn from gamma distributions parametrized by the data. Specifically, we drew 1000 samples from a gamma distribution Γ(*N* ∗ *α*, *θ*), where *α* = *μ*^2^/*σ*^2^, *θ* = *σ*^2^/*μ*, and N is the assumed number of identical neurons to which the ideal observer has access (Britten et al., 1992; Stüttgen and Schwarz, 2008; Wang et al., 2010). We report results using N = 10, but results were qualitatively similar for N = 1, 5, and 20. For each feature velocity, we calculated the true- and false-positive rates for 30 evenly-spaced threshold values between 0 and the maximum response amplitude. We then generated the receiver operator characteristic (ROC) curves by plotting the set of true-positive values vs. the false-positive values. We quantified the theoretical detectability as the area under the ROC curve (AUROC). Because the sensory white noise was presented on adapted trials, it was possible that the adaptive decrease in feature detectability was due in some degree to elevated, white-noise-evoked baseline firing. To isolate the effects of feature response adaptation, we calculated AUROC for a “hybrid” condition using the feature-evoked distributions from adapted trials, and baseline distributions from control trials. Finally, we repeated this analysis by systematically varying the width of the baseline and feature-evoked windows, from 5 ms to 50 ms in 5 ms increments, yielding a time-resolved measure of feature detectability in each condition.

### Feature response latency and synchrony analysis

We sought to estimate the feature response latency of each S1 unit from the PSTH. However, in contrast to the relatively well-populated grand PSTHs, the sparsely-populated PSTHs of individual neurons confound latency calculations. As such, we convolved the spike trains of each neuron with a Gaussian kernel (1 ms standard deviation), yielding a convolved aggregate spike count time series, or a smoothed PSTH. We defined the response latency (T_onset_) for each stimulus condition to be the time at which the smoothed PSTH exceeded a threshold of four standard deviations of the pre-stimulus values (calculated from control trials). We further calculated the T_onset_ adaptation index for each unit as the difference between adapted and control values divided by the sum of the values. Our reported results were qualitatively robust to reasonable choices of onset threshold (2.5, 3.5, 4.0, 5.0, and 6.0 standard deviations) and Gaussian kernel standard deviation (0.5, 1.0, 2.0, and 3.0 ms, not shown).

We calculated the population synchrony of feature responses using the population grand cross-correlogram (CCG) of single-unit spiking, separately for VPm and S1. First, we treated each sensory-responsive unit as a “reference” unit, and calculated the spike times of all other simultaneously-recorded units relative to those of the reference unit. Specifically, for each trial and spike in a reference unit, we calculated the spike times of all other simultaneously-recorded units relative to the reference spike time in a +/− 20 ms window around the reference spike time, and appended these to a grand set of relative spike times. We repeated this for all spikes, trials, and reference units. We then binned the grand set of relative spike times (using 1 ms bins) to generate the grand CCG. We generated a “randomized” grand CCG by producing a grand set of random relative spike times (+/− 20 ms, drawn from a uniform distribution), equal in length to the actual grand set, followed by binning. We then subtracted the randomized CCG from the actual CCG, such that the value at each time lag indicated the number of events beyond that predicted by a random, uncorrelated process. Finally, we divided the CCG by the number of contributing pairs. We defined the mean synchronous spike count to be the sum of the resulting CCG in a +/− 7.5 ms window (Wang et al., 2010). We calculated confidence intervals by re-sampling the grand set of relative spike times with replacement, and re-calculating the CCG and synchrony as described above. To include a pair of neurons in this analysis, we required at least 20 feature-evoked spikes across all trials for both neurons in both the control and adapted conditions. Our results were qualitatively robust to different choices of minimum spike count and synchrony window (not shown). For each feature level, we further calculated the grand CCG and synchrony for a “hybrid” condition, in which each neuron’s spike times from the control condition were subsampled to match the rate of the adapted condition. For the minority of neurons with larger responses in the adapted condition, we retained all spikes from the control condition.

### S1 membrane potential analysis

We removed action potentials from intracellular voltage recordings by first identifying spike times and interpolating between the values 2.5 ms before and 2.5 ms after the peak of the action potential. To identify spike times in each recording, we first calculated the first time derivative time of the membrane potential at each time step. Spike onsets were defined to be positive crossings of five standard deviations of this time series. For each onset, the spike peak time corresponded to the next time at which the derivative was less than or equal to zero. After interpolation, we low-pass-filtered the resulting time series (100 Hz, 3^rd^ order Butterworth). For each trial, we calculated the amplitude and time-to-peak of the subthreshold response to the (sawtooth sensory or optogenetic terminal light) stimulus, in a 50 ms window following stimulus onset. We first subtracted the minimum voltage value in this window on each trial, and defined the amplitude of the response to be the maximum voltage value, and the time-to-peak was the time between stimulus onset and this peak value. We then calculated the across trial median amplitude and time-to-peak.

### Thalamocortical network model

We constructed a simple model of the thalamocortical network using custom scripts written in Python 3.6.10. All code is freely available upon request. We modeled a single cortical barrel as a clustered network of excitatory and inhibitory single-compartment leaky integrate-and-fire (LIF) neurons, subject to excitatory synaptic inputs from a “barreloid” of VPm neurons, and well as excitatory non-thalamic inputs that were independent across cortical neurons. For each of the “control” and “adapted” conditions, we simulated 50 trials, each lasting 150 ms, with a time-step of 0.05 ms.

We modeled a single VPm barreloid as forty independent trains of tonic and burst spikes, drawn from the empirical VPm PSTHs. The ongoing and evoked rates for each neuron were set to the mean empirical values, and were then multiplied by a rate modulation factor drawn from a skewed gamma distribution (with a shape value of 2.0, a scale value of 1.0, then re-scaled to have a mean value of 1.0), to mimic the broad firing rate distributions of VPm neurons previously reported (Pinto et al., 2000; Bruno and Simons, 2002; Wang et al., 2010; Whitmire et al., 2016). Bursts were modeled as pairs of spikes with 2.5 ms ISI.

Non-zero thalamocortical (TC) synaptic weights were drawn from a broad distribution (a rectified Gaussian distribution with mean 1.0 and standard deviation 2.0), to mimic the reported variability in TC synaptic strengths and/or efficacies (Bruno and Simons, 2002; Gabernet et al., 2005; Bruno and Sakmann, 2006; Cruikshank et al., 2007, 2010; Sermet et al., 2019). Mean initial TC synaptic strengths were the same for excitatory and inhibitory neurons (Sermet et al., 2019), but TC convergence was higher for inhibitory neurons (75% for inhibitory neurons, 50% for excitatory neurons). Finally, VPm neurons with the highest firing rates did not synapse onto excitatory neurons (Bruno and Simons, 2002); for each VPm synapse onto an excitatory neuron, if the rate modulation factor of the VPm neuron was greater than 1.5 standard deviations from 1.0, we set that connection to zero, and replaced it with a connection (of the same strength) to a VPm neuron with rate modulation factor below 1.5. In response to a spike in a given thalamic neuron, all TC synapses from that neuron instantly decayed (by a factor of 0.75), followed by exponential recovery (with time constant 25 ms). We assigned a minimum latency of 2 ms (3 ms) to each TC synapse onto an inhibitory (excitatory) neuron, to reflect the slightly faster arrival of thalamic inputs to L4 FS cells (Cruikshank et al., 2007; Kimura et al., 2010). Each TC synapse was assigned an additional temporal latency value between 0 and 1 ms, drawn from a uniform distribution, which helped to promote evoked cortical firing that was less time-locked across the network. Importantly, the higher TC convergence for inhibitory neurons, the rate-dependent TC connectivity described above, the shorter TC synaptic latencies onto inhibitory neurons, and intrinsic neuronal and intracortical connection parameters (see below) together supported strong feedforward inhibition in this model network, despite the equivalent mean TC synaptic weights onto the two neuron types.

Each thalamic spike resulted in a postsynaptic conductance waveform in each postsynaptic cortical neuron, with excitatory reversal potential 50 mV. Given a thalamic spike at time zero, we modeled the resulting postsynaptic conductance waveform *g*(*t*) as a conductance amplitude *ḡ* multiplied by a gating variable *G*(*t*). The gating variable at a given TC synapse had the form of a difference of two exponentials, resulting in rise and subsequent decay in the synaptic conductance following a presynaptic spike at time zero:

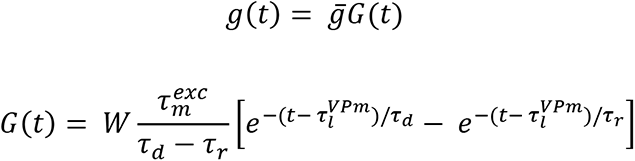

where W is the weight of the TC synapse, 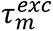 is the membrane time constant of excitatory neurons, *τ*_*d*_ is the decay time constant, *τ*_*r*_ is the rise time constant, and 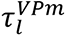 is the latency of the synapse. We selected rise and decay time constants of 0.2 and 1.0 ms, respectively, for all TC synapses, yielding a relatively fast rise and slow decay. We include 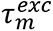 in the numerator of the leading factor so that the gating variable is unitless. Note that this means the across-synapse mean integrated value of G(t) will be equal to 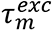, which should be taken into account when interpreting absolute values of TC synaptic conductance amplitudes. We tuned the TC synaptic conductance amplitude (to 30 nS) such that near-simultaneous firing of multiple thalamic neurons was required to evoke action potentials in target neurons (Bruno and Sakmann, 2006), when intracortical synaptic strengths were set to zero.

Each cortical network neuron also received random excitatory “nonthalamic” inputs, to promote baseline network activity. For each neuron, spike times were drawn from a homogeneous Poisson process, with rate 60 Hz. The synaptic weights for random external inputs were drawn from a rectified, normalized Gaussian distribution (with mean of 1.0, standard deviation of 2.0, then rectified, then divided by its mean), to support variable ongoing firing rates across network neurons. Each random spike resulted in a postsynaptic conductance waveform, with the same rise and decay time constants as for thalamocortical inputs, and with conductance amplitude 60 nS.

We modeled a single cortical column as a network of 800 excitatory and 100 inhibitory neurons, which approximates the relative numbers of these neuron subtypes across layers in S1 (Lefort et al., 2009). Excitatory-to-excitatory connection probability was 4%, excitatory-to-inhibitory was 20%, inhibitory-to-excitatory was 17.5%, and inhibitory-to-inhibitory was 17.5%. While these numbers are approximately three times lower than what has been reported within a column in S1 (Avermann et al., 2012), these lower connection probabilities were necessary to maintain network stability, and the relative values are in qualitative agreement with experimental measurements. We imposed spatial clustering of connectivity via a “small-world” network connectivity (Litwin-Kumar and Doiron, 2012; Wright et al., 2017a, 2017b; Hoseini et al., 2019), with 10% re-wiring probability. First, all neurons were organized on a continuous ring. Then, outputs from a given cortical excitatory network neuron were projected to 32 of its nearest excitatory neighbors. For each of these outputs, the connection was reassigned to another randomly-selected (not previously connected) excitatory neuron, with probability 10%. In other words, most connections were nearest-neighbor, but some were random (and possibly long-distance). Each nonzero connection weight was drawn from a heavy-tailed beta distribution (with alpha value 0.11, beta value 1.0, and then re-scaled to have mean value 1.0), to approximate the variable intracortical synaptic strengths previously reported in cortex (Avermann et al., 2012; Cossell et al., 2015; Pala and Petersen, 2015). This process was repeated for all neuron pair types, according to the connection probabilities listed above. All intracortical synapses were assigned a latency of 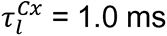, and rise and decay time constants of 0.2 and 1.0 ms, respectively, and the synaptic gating variables evolved as described above for TC synapses. Intracortical synaptic conductance amplitudes were 27.5 nS for excitatory-to-excitatory connections, 30 nS for excitatory-to-inhibitory connections, 40 nS for inhibitory-to-excitatory connections, and 30 nS for inhibitory-to-inhibitory connections.

For network LIF neurons, we selected intrinsic neuronal properties that were consistent with previous modeling studies, and/or were motivated by previous experimental work. We set the membrane time constant of excitatory neurons (inhibitory neurons) to be 30 ms (20 ms), reflecting the moderately higher input resistances of cortical excitatory neurons reported in awake mouse (Gentet et al., 2010). Inhibitory neurons also had shorter refractory periods (1 ms vs. 2 ms), which supported higher firing rates, as observed previously (Bruno and Simons, 2002; Khatri et al., 2004; Gentet et al., 2010; Taub et al., 2013). Excitatory neurons were subject to an inhibitory spike-rate adaptation conductance, with a 1 ms rise time constant, and 30 ms decay time constant. We tuned the time constants and amplitude of this conductance such that the ISI of spikes evoked by tonic current injection gradually increased over the duration of the stimulus. This conductance served to stabilize excitatory firing, and therefore the network as a whole. For each neuron, the leak reversal potential *E*_*leak*_ was drawn from a uniform distribution between −70 mV and −60 mV (which added variability in spike timing across the network), and the leak conductance amplitude was 100 nS. All neurons had a spike threshold of − 45 mV, reset to −59 mV immediately after a spike, and were held at the reset value during the refractory period.

For each condition, we simulated 50 trials, each lasting 150 ms (including a 50 ms “buffer window” to allow the network to reach stead-state, a 50 ms “pre-stimulus” window, and a 50 ms “post-stimulus” window), with a time-step of 0.05 ms. At each time-step, the membrane potential V of a given network neuron of type *Y* (where *Y* is either *E* or *I*, representing an excitatory or inhibitory neuron) evolved according to the equations

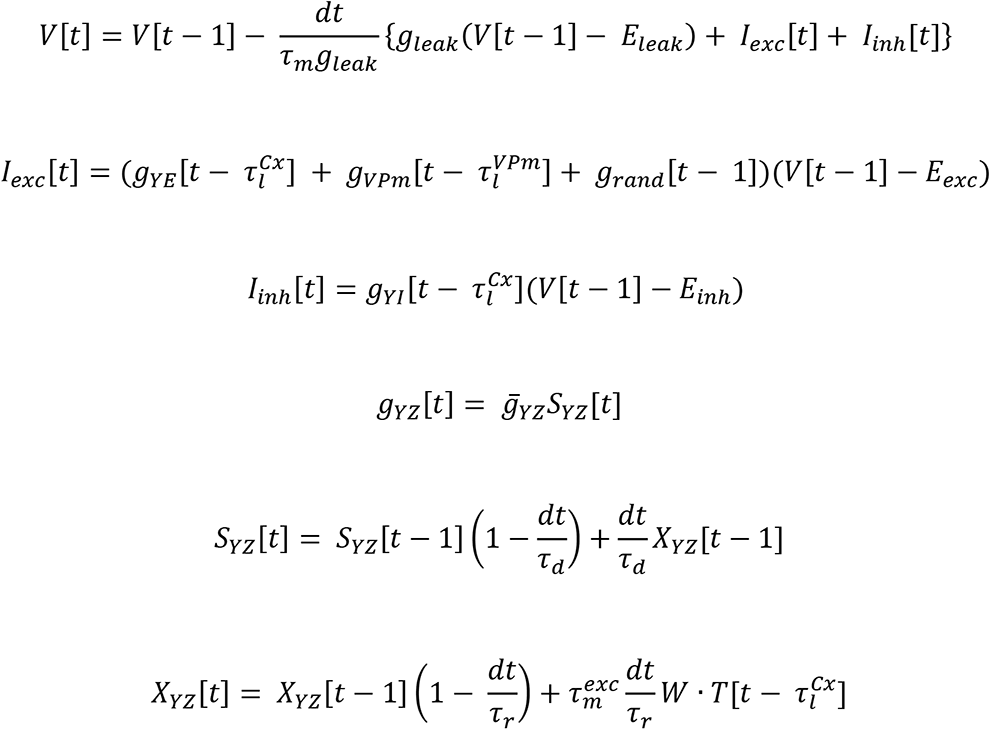

where *g*_*YZ*_ is the summed synaptic conductance time series from all presynaptic network neurons of type *Z = E|I* onto this target neuron of type *Y = E|I*, *g_VPm_* is the summed synaptic conductance time series from all presynaptic VPm neurons, and *g*_*rand*_ is an excitatory synaptic conductance resulting from Poisson process spike times that were drawn independently for each neuron. The equations for *S*_*YZ*_ and *X*_*YZ*_ represent the implementation of the gating variable *G(t)* described above, associated with conductance of type *Z* onto neuron of type *Y*. *W* is the (input) synaptic weight matrix for presynaptic neurons of type *Z* onto neurons of type *Y*, and *T* is the binary spike train sequence for all presynaptic neurons of type *Z*. Excitatory neurons were also subject to an inhibitory spike-rate-adaptation current, described above.

We quantified model responses by calculating the peaks of the grand PSTHs for excitatory and inhibitory LIF neurons and divided “adapted” values by “control” values, yielding the normalized adapted response. We generated grand cross-correlograms (as described above) for 200 randomly-selected excitatory-excitatory and inhibitory-inhibitory pairs, and for 100 VPm-VPm pairs.

We further employed alternate models to parse the roles played by synchronous thalamic spikes and feedforward inhibition. For the “reduced synch” models, we maintained the mean spike rates of the original model, but manually adjusted drawn VPm spike times to reduce synchrony. Specifically, if a drawn VPm spike time was within +/− 5 ms of the empirical PSTH peak time, we shifted the spike to a random later time with probability P(shift), within approximately 20 ms of the PSTH peak. We repeated this process for P(shift) = 0.1, 0.2, 0.3, and 0.4. For the “Identical TC Connectivity” model, excitatory and inhibitory neurons had the same TC convergence values (50%), TC synaptic latencies were identical for excitatory and inhibitory neurons,s and we did not require that VPm neurons with the highest rates synapse exclusively onto inhibitory neurons.

## Results

To investigate the adaptive effects of persistent sensory stimulation on S1 sensory responses during wakefulness, we presented precise deflections to a single whisker of the awake, head-fixed mouse using a computer controlled galvanometer, and recorded extracellular spiking activity in the corresponding principal column of S1, and/or principal barreloid of VPm (Fig. 1A, see Methods). We presented punctate “sawtooth” sensory features either in isolation or embedded in an adapting background stimulus (frozen sensory white noise, Fig. 1A). The punctate stimulus feature captures the basic nature of the high velocity “stick-slip” whisker motion events that occur as a result of whisker contacts with larger surface irregularities during active sensation (Ritt et al., 2008; Wolfe et al., 2008; Jadhav et al., 2009; Jadhav and Feldman, 2010). During whisker contacts with surfaces, these stick-slip events are embedded in patterns of smaller-amplitude, irregular deflections (Jadhav and Feldman, 2010), simplistically captured here utilizing low-amplitude, repeatable background white noise whisker stimulation, which has been shown to alter perceptual reports in behaving rats (Waiblinger et al., 2015, 2018) and adapt thalamic feature responses under anesthesia (Whitmire et al., 2016). We first characterized the effects of the background stimulus on baseline and feature-evoked cortical firing during wakefulness (Figs. 1 – 3), and then sought to identify the mechanisms underlying these effects through a battery of additional experiments (Figs. 4 – 7) and thalamocortical network modeling (Fig. 8).

**Figure 1.**
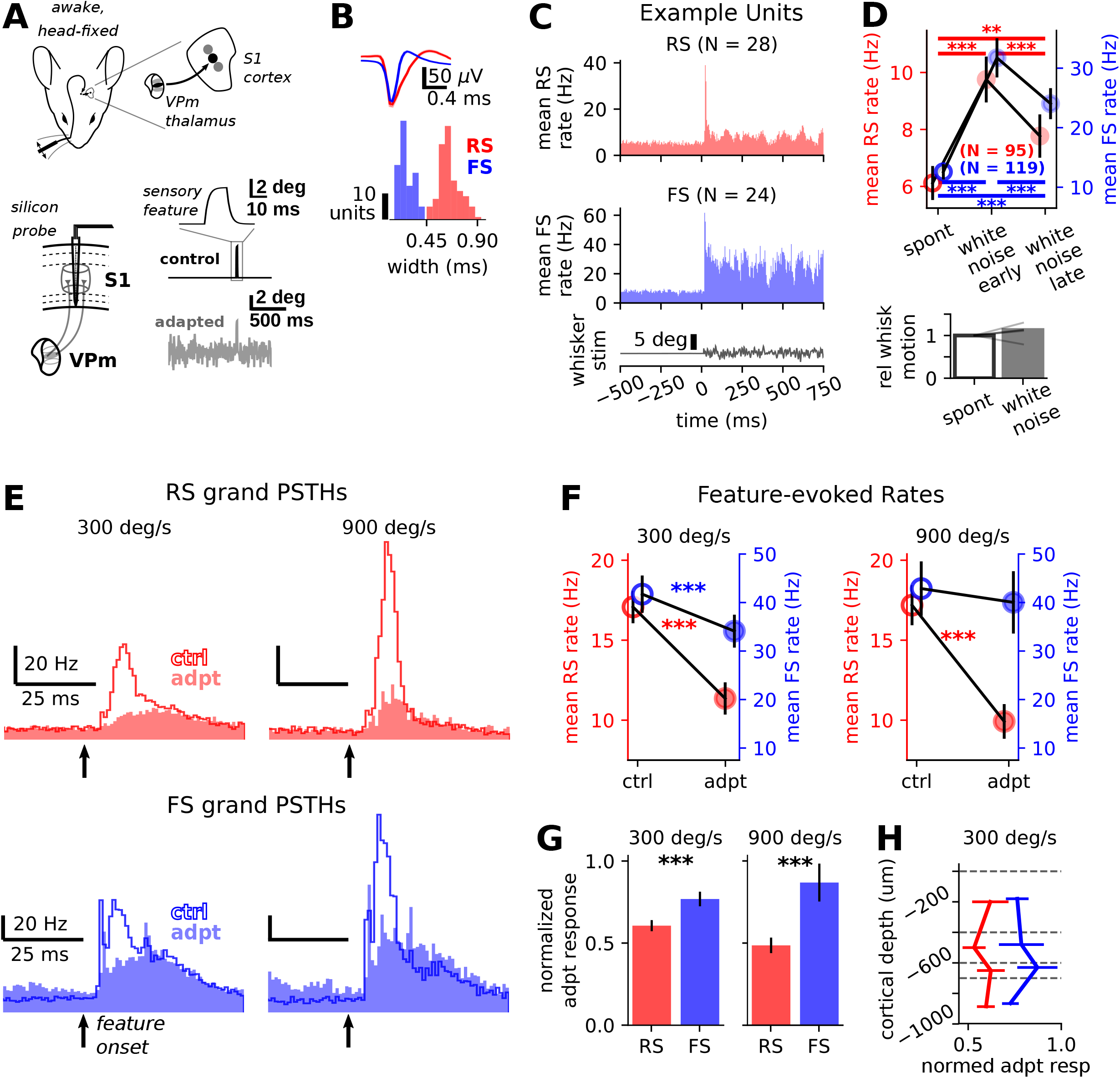
S1 exhibits sensory adaptation during wakefulness, and regular-spiking neurons are more profoundly adapted than fast-spiking neurons. **A.** Experimental setup. We recorded in S1 of the awake, head-fixed mouse while presenting precise single-whisker stimulation. “Sawtooth” punctate sensory features were delivered either in isolation (control condition) or embedded in sensory white noise (adapted condition). Top: Grand mean +/− SEM waveforms for all well-isolated fast-spiking (FS, blue, N = 95) and regular-spiking (RS, red, N = 119) significantly responsive single-units recorded in S1 of awake mice (see Methods). Bottom: distribution of mean waveform widths (time from trough to subsequent peak) for all units, with color denoting RS and FS designation. Grand PSTHs for regular-spiking (RS, putative excitatory, top) and fast-spiking (FS, putative inhibitory, bottom) from a subset of all recording sessions (for which we used the same white noise stimulus). D. Top: grand-average mean (+/− SEM) rates for spontaneous activity (i.e., no sensory stimulation) and “early” (0 – 200 ms) and “late” (500 700 ms) windows following onset of sensory white noise (with *** indicating p < 0.0005, Wilcoxon signed-rank test). RS: spontaneous rate = 6.12 +/− 0.60 Hz; white noise early rate = 9.75 +/− 0.80 Hz; white noise late rate = 7.77 +/− 0.76 Hz, mean +/− SEM. Spont vs. white noise early: W = 274, p = 4.59 × 10^−10^; spont vs. white noise late: W = 799.5, p = 5.92 × 10^−4^; white noise early vs. white noise late: W = 425, p = 9.95 × 10^−9^, Wilcoxon signed-rank test, N = 119 units from 19 recording sessions, FS: spontaneous rate = 12.59 +/− 1.29 Hz, white noise early rate = 31.73 +/− 3.29 Hz; white noise late rate = 24.04 +/− 2.6 Hz, mean +/− SEM. Spont vs. white noise early: W = 4.0, p = 9.13 × 10^−15^; spont vs. white noise late: W = 237.5, p = 3.34 × 10^−11^; white noise early vs. white noise late: W = 220.0, p = 1.88 × 10^−11^, Wilcoxon signed-rank test, N = 95 units from 19 sessions. Bottom: relative whisker motion in the absence (“spont”) and presence (“white noise”) of white noise whisker stimulation, from five recording sessions with simultaneous electrophysiology and whisker videography (see Methods). E. Grand PSTHs for all responsive RS (top, N = 119) and FS (bottom, N = 95) units, for two punctate stimulus velocities. F. Across-neuron mean (+/− SEM) firing rates for all responsive neurons, for 300 deg/s (left) and 900 deg/s (right) punctate stimuli (*: 0.01 ≤ p < 0.05; ***: p < 0.001, Wilcoxon singed-rank test). RS 300 deg/s mean +/− SEM control: 17.08 +/− 1.02, adapted: 11.36 +/− 1.01, 33.5% decrease, W = 523.0, p = 1.03 × 10^−15^, Wilcoxon signed-rank test, N = 119 units from 19 recording sessions; 900 deg/s control: 17.19 +/− 1.25 Hz, adapted: 9.92 +/− 1.08 Hz, 42.3% decrease, W = 43.0, p = 1.47 × 10^−8^, N = 49 units from 8 sessions; FS: 300 deg/s control: 41.72 +/− 3.85 Hz, adapted: 34.1 +/− 3.39 Hz, 18.3% decrease, W = 785.5, p = 2.90 × 10^−8^, N = 95 units from 19 sessions; 900 deg/s control: 42.88 +/− 5.61 Hz, adapted: 40.03 +/− 6.43 Hz, 6.6% decrease, W = 156.5, p = 0.19, N = 29 units from 8 sessions. G. Population median (+/− SEM) normalized adapted responses for all responsive RS (red) and FS (blue) neurons (see Methods; ***: p < 0.001, Kruskal-Wallis test). RS 300 deg/s median normed adapted response = 0.61, FS median normed adapted response = 0.77, H = 16.94, p = 3.85 × 10^−5^, Kruskal-Wallis test; 900 deg/s RS median normed adapted response = 0.49, FS median normed adapted response = 0.87, H = 14.91, p = 1.13 × 10^−4^. H. Population median normalized adapted responses (+/− SEM) by binned cortical depth (see Methods).

**Figure 2.**
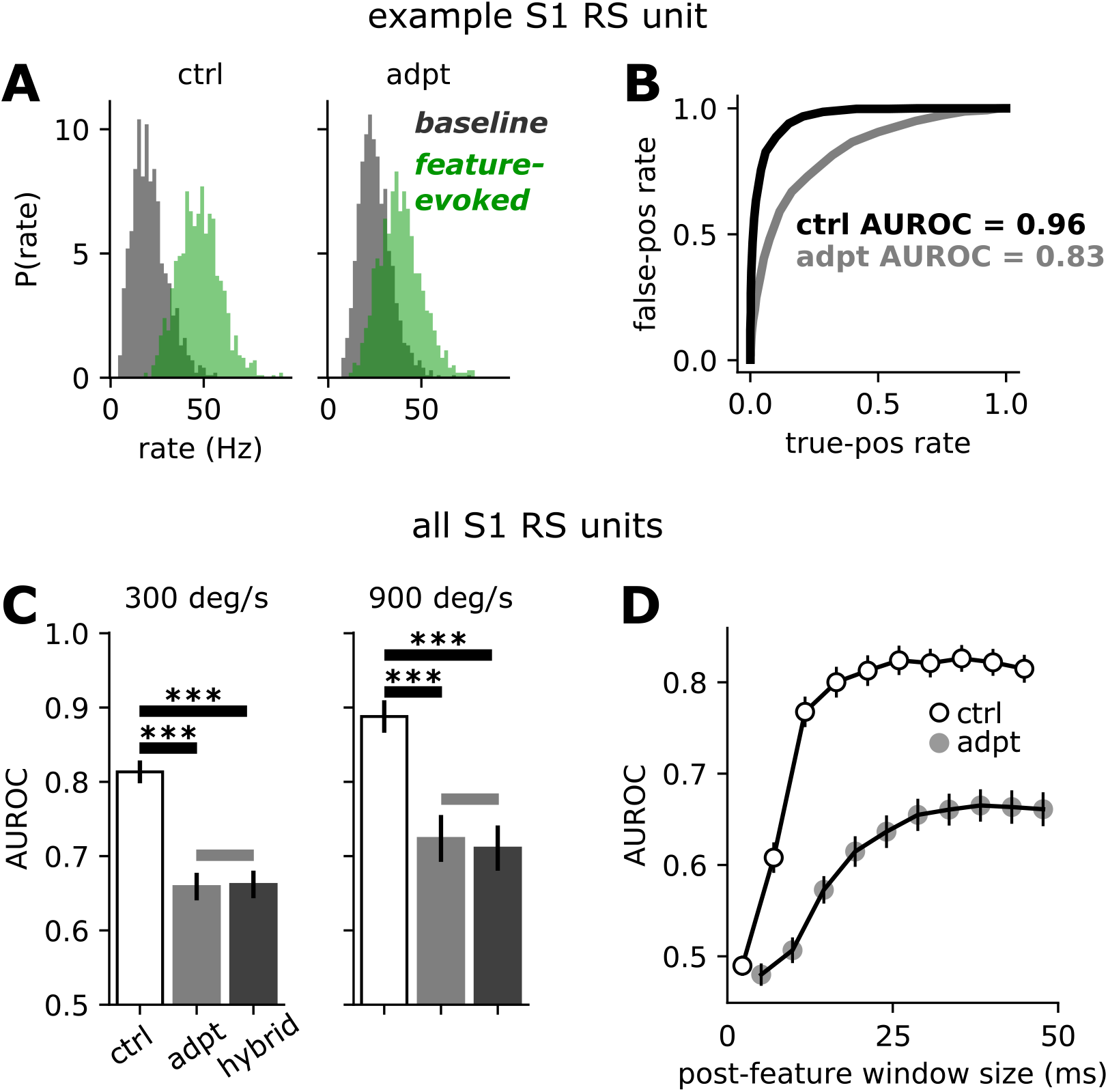
Adaptation reduces the theoretical detectability of punctate sensory stimuli. **A.** Baseline and feature-evoked population spike rate distributions for one example RS unit (see Methods). For this unit (qualitatively representative of the average), adaptation decreased the mean population response rate, increasing the overlap of the baseline and feature-evoked distributions (right). **B.** Receiver operator characteristic (ROC) curves for example unit in (B), for the control (black line) and adapted (gray line) conditions, and associated area under ROC curve (AUROC) values. **C.** Grand average mean (+/− SEM) theoretical detectability (AUROC) vs. stimulus condition for all significantly responsive RS units (***: p < 0.0005, Wilcoxon signed-rank test). 300 deg/s control: mean +/− SEM AUROC = 0.81 +/− 0.02, adapted: 0.66 +/− 0.02, hybrid: 0.66 +/− 0.02, control vs. adapted: W = 1936.0, p = 2.09 × 10^−11^; control vs. hybrid: W = 1154.0, p = 1.91 × 10^−16^; adapted vs. hybrid: W = 5243.0, p = 0.81, Wilcoxon signed-rank test, N = 119 units from 19 recording sessions; 900 deg/s control: mean +/− SEM AUROC = 0.89 +/− 0.02, adapted: 0.73 +/− 0.03, hybrid: 0.71 +/− 0.03; control vs. adapted: W = 166.0, p = 3.18 × 10^−6^; control vs. hybrid: W = 106, p = 1.78 × 10^−7^; adapted vs. hybrid: W = 528.0, p = 0.21, N = 49 units from 8 recording sessions.

**Figure 3.**
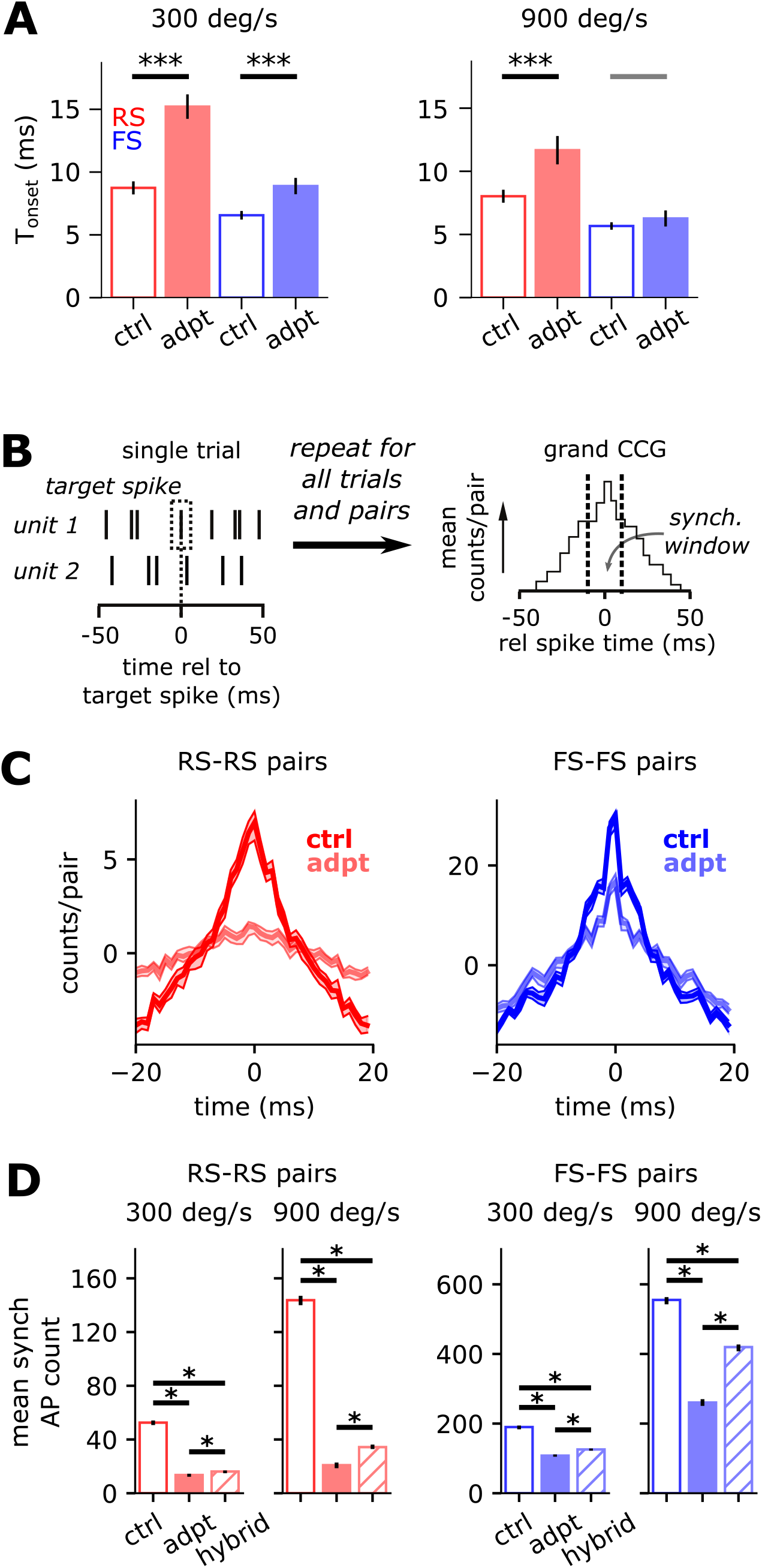
Adaptation increases response latency and reduces pairwise synchronous spiking in S1. **A.** Grand mean onset times for RS (red) and FS (blue) neurons, for control and adapted responses to 300 deg/s stimulus (***: p < 0.001; single gray bar: p ≥ 0.05, Wilcoxon signed-rank test, control vs. adapted). 300 deg/s RS mean +/ SEM control: T_onset_ = 8.81 +/− 0.53 ms, adapted: 15.31 +/− 0.98 ms, 73.9% increase, W = 397.0, p = 3.11 × 10^−11^, Wilcoxon signed-rank test, N = 119 units; FS control: T_onset_ = 6.69 +/− 0.37 ms, adapted: 8.95 +/− 0.7 ms, 33.9% increase, W = 245.0, p = 8.11 × 10^−7^, N = 95 units; 900 deg/s RS control: T_onset_ = 7.89 +/− 0.49 ms, adapted: 11.36 +/− 1.07 ms, 43.9% increase, W = 90.0, p = 1.70 × 10^−5^, N = 49 units; FS control: T_onset_ = 5.67 +/− 0.29 ms, adapted: 6.27 +/− 0.63 ms, 10.5% increase, W = 20.5, p = 0.27, N = 29 units. **B.** Illustration of synchronous spike-count calculation. The grand cross-correlogram (CCG) was constructed using all valid pairs of simultaneously-recorded S1 units, then scaled by the number of contributing pairs, and shuffle-corrected (see Methods). The synchronous spike count was the number of spikes in a +/− 7.5 ms window around zero lag. **C.** Grand RS-RS (left) and FS-FS (right) CCGs for responses to 300 deg/s stimulus, for the control (dark lines) and adapted (light lines) conditions. Bands indicate 97.5% confidence intervals (from re-sampling spikes with replacement, see Methods). **D.** Synchronous AP counts for control, adapted, and “hybrid” conditions (see Methods), calculated from grand CCGs. (Error bars indicate 97.5% confidence intervals; *: p < 0.025, re-sampling spikes with replacement, see Methods.) RS-RS pairs: 300 deg/s mean +/− 97.5% confidence interval control: synch AP count = 54.15 +/− 3.58 spikes/pair; adapted: 13.69 +/− 2.05 spikes/pair; hybrid: 16.61 +/− 1.67 spikes/pair; N = 189 valid pairs from 17 sessions; 900 deg/s control: 141.93 +/− 5.42 spikes/pair; adapted: 21.18 +/− 2.8 spikes/pair; hybrid: 34.87 +/− 3.84 spikes/pair; N = 55 pairs from 6 sessions. FS-FS pairs: 300 deg/s control: 191.03 +/− 7.64 spikes/pair; adapted: 106.54 +/− 7.49 spikes/pair; hybrid: 126.33 +/− 6.69 spikes/pair; N = 118 pairs from 17 sessions; 900 deg/s control: 558.27 +/− 26.03 spikes/pair; adapted: 257.15 +/− 19.67 spikes/pair; hybrid: 423.61 +/− 17.97 spikes/pair; N = 33 pairs from 6 sessions. See Methods for definition of “valid pairs”.

**Figure 4.**
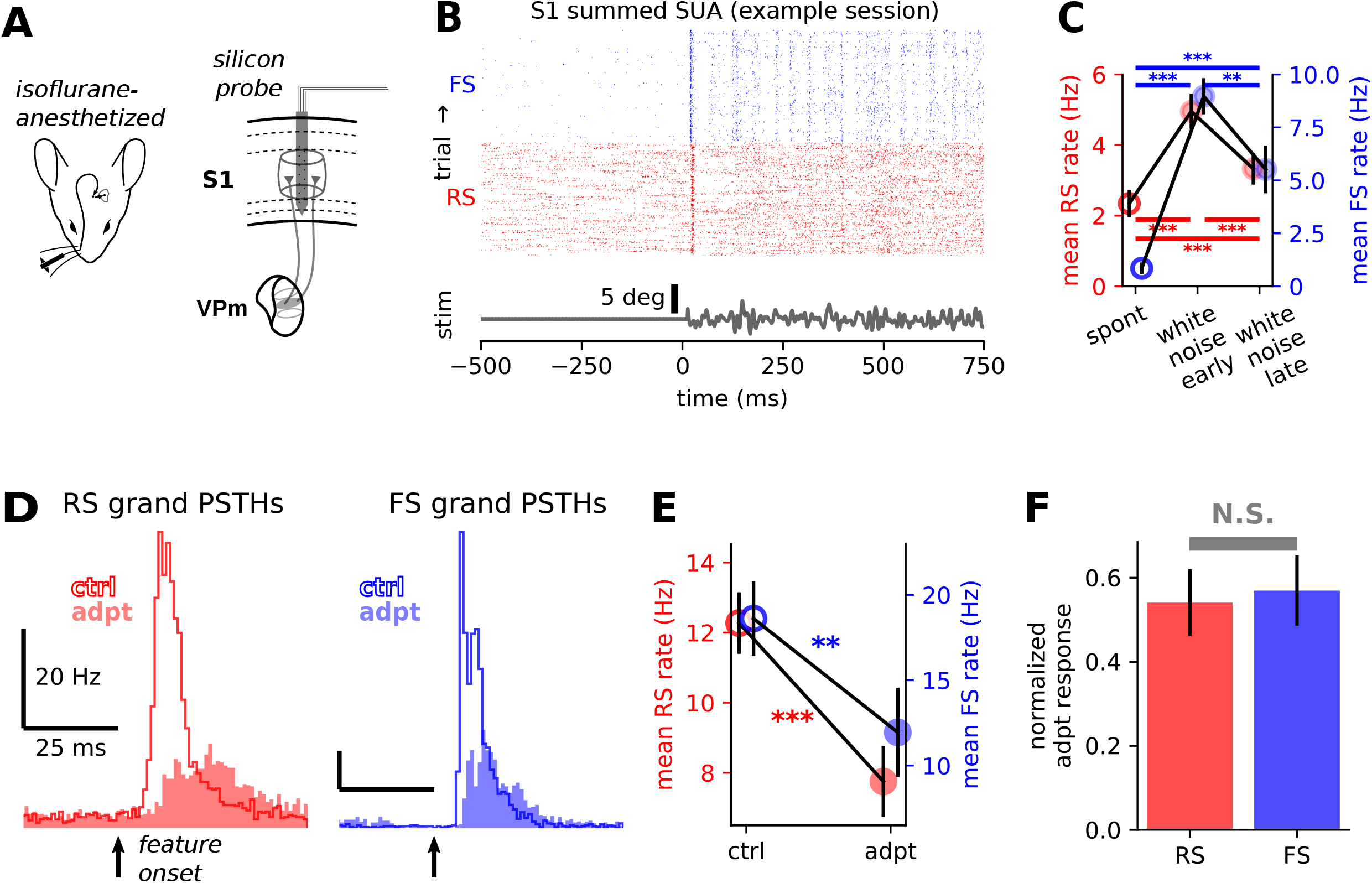
S1 sensory responses are also adapted under anesthesia, but RS cells are not differentially adapted. **A.** Experimental setup. S1 sensory responses were recorded in mice lightly-anesthetized with isoflurane (see Methods). **B.** Summed spiking activity of regular-spiking (RS, putative excitatory) and fast-spiking (FS, putative inhibitory) from one example recording session. Each row indicates spike times of all simultaneously-recorded RS (red) and FS (blue) neurons on a single trial. **C.** Grand-average mean (+/− SEM) rates for spontaneous activity (i.e., no sensory stimulation) and “early” (0 – 200 ms) and “late” (500 – 700 ms) windows following onset of sensory white noise (with *** indicating p < 0.0005, Wilcoxon signed-rank test). RS: spontaneous rate = 2.34 +/− 0.38 Hz; white noise early rate = 4.96 +/− 0.50 Hz; white noise late rate = 3.33 +/− 0.45 Hz, mean +/− SEM. Spont vs. white noise early: W = 55, p = 1.13 × 10^−7^; spont vs. white noise late: W = 191, p = 3.89 × 10^−4^; white noise early vs. white noise late: W = 214, p = 3.61 × 10^−4^, Wilcoxon signed-rank test, N = 46 units from 14 recording sessions, FS: spontaneous rate = 0.85 +/− 0.28 Hz, white noise early rate = 8.97 +/− 0.85 Hz; white noise late rate = 5.52 +/− 1.13 Hz, mean +/− SEM. Spont vs. white noise early: W = 0, p = 5.95 × 10^−5^; spont vs. white noise late: W = 6, p = 1.41 × 10^−4^; white noise early vs. white noise late: W = 20, p = 9.02 × 10^−4^, Wilcoxon signed-rank test, N = 21 units from 14 sessions. **D.** Sensory response grand PSTHs for all responsive RS (left, N = 46) and FS (right, N = 21) units recorded under anesthesia, for 300 deg/s punctate stimulus velocity. **E.** Grand mean (+/− SEM) rates for cells contributing to PSTHs in (D). RS mean +/− SEM control: 12.27 +/− 0.88 Hz, adapted: 7.75 +/− 1.01, 36.8% decrease, W = 184, p = 9.82 × 10^−5^; FS control: 18.61 +/− 2.19 Hz, adapted: 11.94 +/− 2.61, 35.8% decrease, W = 21, p = 0.001. **F.** Population median (+/− SEM) normalized adapted responses for all responsive RS (red) and FS (blue) neurons (see Methods). RS 300 deg/s median normed adapted response = 0.54, FS median normed adapted response = 0.57, H = 0.01, p = 0.91, Kruskal-Wallis test.

**Figure 5.**
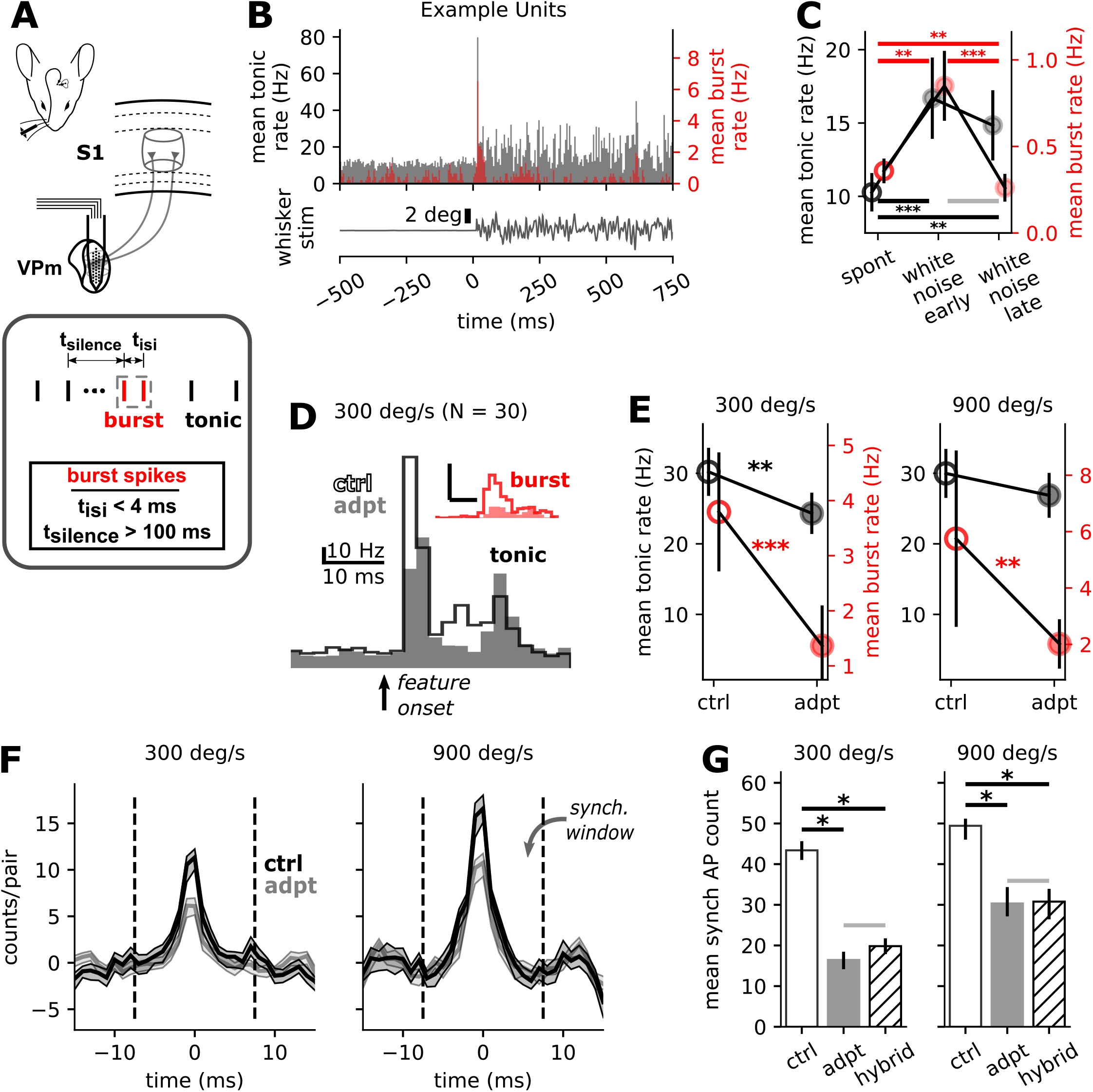
Adaptation reduces tonic and burst firing rates, and synchronous spike counts, in VPm sensory responses. **A.** Top: experimental setup. We recorded extracellular spiking in VPm of the awake mouse, primarily using high-density silicon probes (see Methods). Bottom: criteria for classification of putative tonic (black) and burst (red) VPm spikes. **B.** Grand PSTHs for putative tonic (black) and burst (red) VPm spikes from a subset of all recording sessions. **C.** Grand-average mean (+/− SEM) rates for spontaneous activity (i.e., no sensory stimulation) and “early” (0 – 200 ms) and “late” (500 700 ms) windows following onset of sensory white noise (with *** indicating p < 0.0005, Wilcoxon signed-rank test). Tonic: spontaneous rate = 10.28 +/− 1.31 Hz; white noise early rate = 16.69 +/− 2.77 Hz; white noise late rate = 14.84 +/− 2.39 Hz, mean +/− SEM. Spont vs. white noise early: W = 42, p = 8.91 × 10^−5^; spont vs. white noise late: W = 75, p = 1.12 × 10^−3^; white noise early vs. white noise late: W = 135, p = 0.045, Wilcoxon signed-rank test; burst: spontaneous rate = 0.36 +/− 0.07 Hz, white noise early rate = 0.85 +/− 0.2 Hz; white noise late rate = 0.26 +/− 0.08 Hz, mean +/− SEM. Spont vs. white noise early: W = 65.5, p = 3.0 × 10^−3^; spont vs. white noise late: W = 51, p = 4.65 × 10^−3^; white noise early vs. white noise late: W = 37, p = 4.35 × 10^−4^, Wilcoxon signed-rank test, N = 30 units from 9 sessions. **D.** Grand PSTHs for tonic (black) and burst (red) spikes from all putative VPm neurons, for 300 deg/s punctate stimulus. Note the presence of both a short-latency primary peak, and a shorter, secondary peak in tonic firing rates (Fig. 5E). This secondary peak in the grand PSTH resulted from a subset of neurons with both early and late responses – often within individual trials – and was likely evoked by the return of the whisker to resting position in the second half of the sawtooth waveform. **E.** Across-neuron mean (+/− SEM) firing rates for all putative VPm neurons (**: 0.001 ≤ p < 0.01; ***: p < 0.001, Wilcoxon singed-rank test). Tonic 300 deg/s mean +/− SEM rate control: 30.2 +/− 3.41 Hz, adapted: 24.32 +/− 2.92 Hz, 19.5% decrease, W = 83, p = 0.002, Wilcoxon signed-rank test, N = 30 units; 900 control: 29.98 +/− 3.47 Hz, adapted: 26.87 +/− 3.18 Hz, W = 44.5, p = 0.22, N = 16 units; burst: 300 deg/s control: 3.79 +/− 1.08, adapted: 1.37 +/− 0.73 Hz, 64.0% decrease, W = 47, p = 6.21 × 10^−4^; 900 deg/s control: 5.75 +/− 3.13 Hz, adapted: 2.02 +/− 0.88, 64.9% decrease, W = 5, p = 4.6 × 10^−3^. **F.** Grand shuffle-corrected cross-correlograms for all simultaneously-recorded putative VPm neurons, for the control (black) and adapted (gray) conditions. Bands indicate 97.5% confidence intervals (from re-sampling spikes with replacement, see Methods). **G.** Synchronous AP counts for control, adapted, and “hybrid” conditions (see Methods), calculated from grand CCGs. (Error bars indicate 97.5% confidence intervals; *: p < 0.025, re-sampling spikes with replacement, see Methods.) 300 deg/s mean +/− 97.5% confidence interval control: synch AP count = 46.69 +/− 8.12 spikes/pair; adapted: 17.29 +/− 4.0 spikes/pair; hybrid: 20.4 +/− 4.25 spikes/pair; N = 48 valid pairs from 7 sessions; 900 deg/s control: 51.38 +/− 8.22 spikes/pair; adapted: 30.81 +/− 6.3 spikes/pair; hybrid: 31.3 +/− 5.35 spikes/pair; N = 37 pairs from 3 sessions. (See Methods for definition of “valid pairs”.)

**Figure 6.**
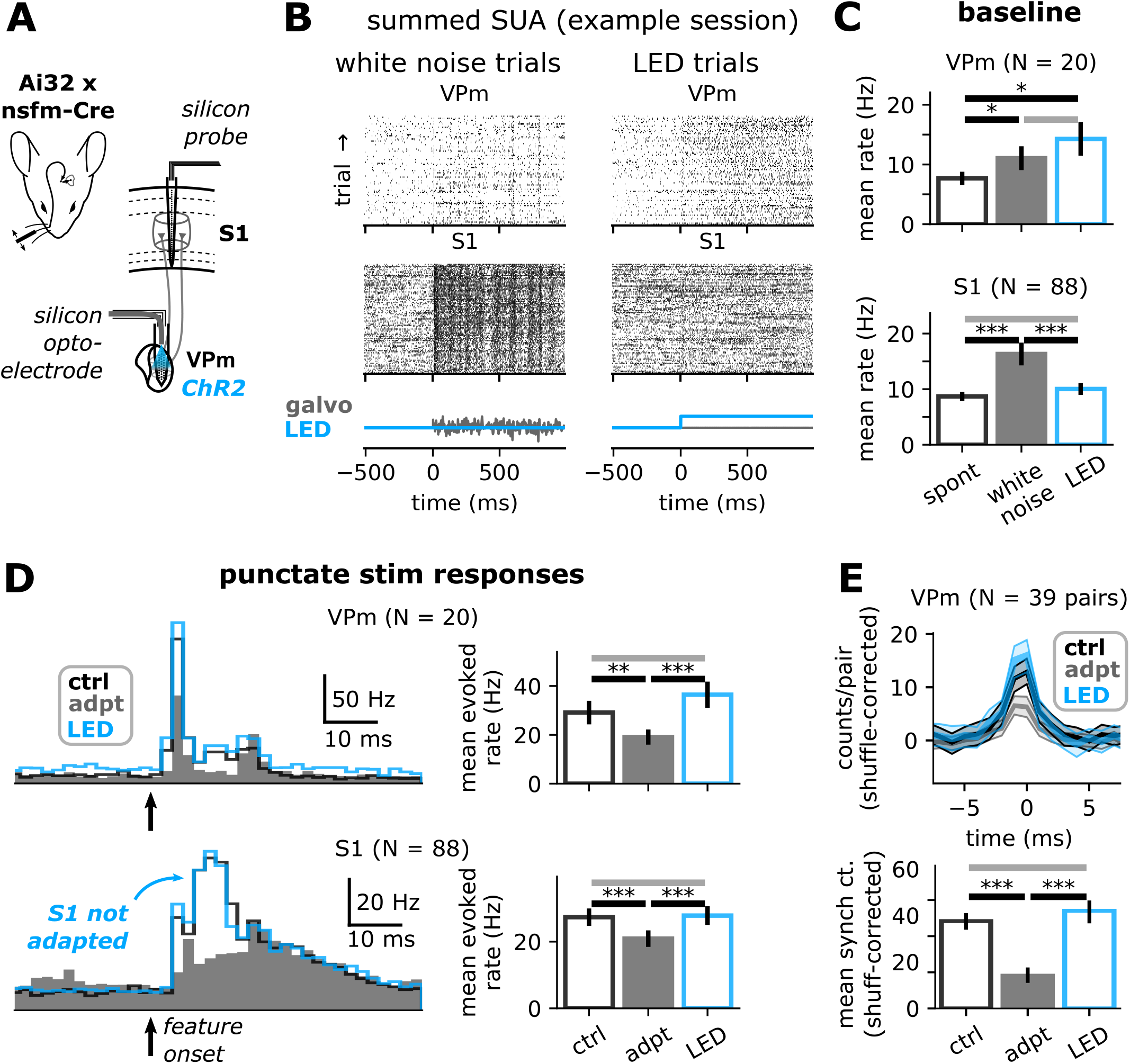
Optogenetic elevation of baseline VPm firing rate does not adapt S1 sensory responses. **A.** Experimental setup. We recorded extracellular spiking activity from topographically-aligned VPm barreloids and S1 barrels in awake, head-fixed, transgenic mice expressing Channelrhodopsin in VPm/VPl neurons (see Methods). **B.** Summed spiking activity of all well-isolated, responsive putative VPm (top) and S1 (bottom) units from one example recording session. Each row indicates spike times of all such simultaneously-recorded units in that brain region on a single trial. **C.** Grand mean (+/− SEM) firing rates for spontaneous activity (“spont”), and during presentation of the adapting sensory stimulus (“white noise”) and optogenetic depolarization of VPm (“LED”), for VPm (top) and S1 (bottom; *: 0.005 ≤ p < 0.025; **: 5 × 10^−4^ ≤ p < 5 × 10^−3^; ***: p < 5 × 10^−4^). VPm mean +/− SEM rate spontaneous: 7.68 +/− 1.1 Hz, white noise: 11.05 +/− 1.99 Hz, LED: 14.27 +/− 2.81 Hz; spont vs. white noise, W = 37, p = 0.011, Wilcoxon signed-rank test; spont vs. LED: W = 37, p = 0.011; white noise vs. LED: W = 102, p = 0.91, N = 20 units from 6 sessions; S1 spontaneous: 8.87 +/− 0.82 Hz, white noise: 16.32 +/− 2.05 Hz, LED: 10.03 +/− 1.05 Hz; spont vs. white noise: W = 445, p = 5.05 × 10^−10^; spont vs. LED: W = 1568.5, p = 0.11; white noise vs. LED: W = 1013, p = 8.42 × 10^−5^, N = 88 units from six sessions. **D.** Left: grand PSTHs for each stimulus condition, for VPm (top) and S1 (bottom). Right: Grand mean (+/− SEM) firing rates for each condition. Asterisks as in (C). VPm mean +/− SEM rate control: 29.12 +/− 4.82 Hz, adapted: 19.07 +/− 3.1 Hz, LED: 36.44 +/− 5.35 Hz; control vs. adapted: W = 26, p = 0.003; adapted vs. LED: W = 11, p = 4.5 × 10^−4^; control vs. LED: W = 55, p = 0.06. S1 control: 27.33 +/− 2.61 Hz, adapted: 20.91 +/− 2.47 Hz, LED: 27.82 +/− 2.79 Hz; control vs. adapted: W = 502.5, p = 1.39 × 10^−9^; adapted vs. LED: W = 424, p = 2.86 × 10^−10^; control vs. LED: W = 1837, p = 0.89, Wilcoxon signed-rank test. **E.** Top: Grand VPm CCGs for each stimulus condition. Bands indicate 99.9% confidence intervals (re-sampling spikes with replacement (see Methods). Bottom: synchronous spike counts calculated from CCGs in (E, Top), for each stimulus condition. Error bars indicate 99.9% confidence intervals (re-sampling spikes with replacement (see Methods). Mean +/− 99.9% confidence interval synch AP count control: 47.14 +/− 11.0 spikes/pair, LED: 51.83 +/− 12.94 spikes/pair, adapted: 20.08 +/− 8.53 spikes/pair, p ≥ 0.05 control vs. LED, p < 0.001 control vs. adapted, re-sampling spikes with replacement, N = 36 pairs from 19 units in five sessions.

**Figure 7.**
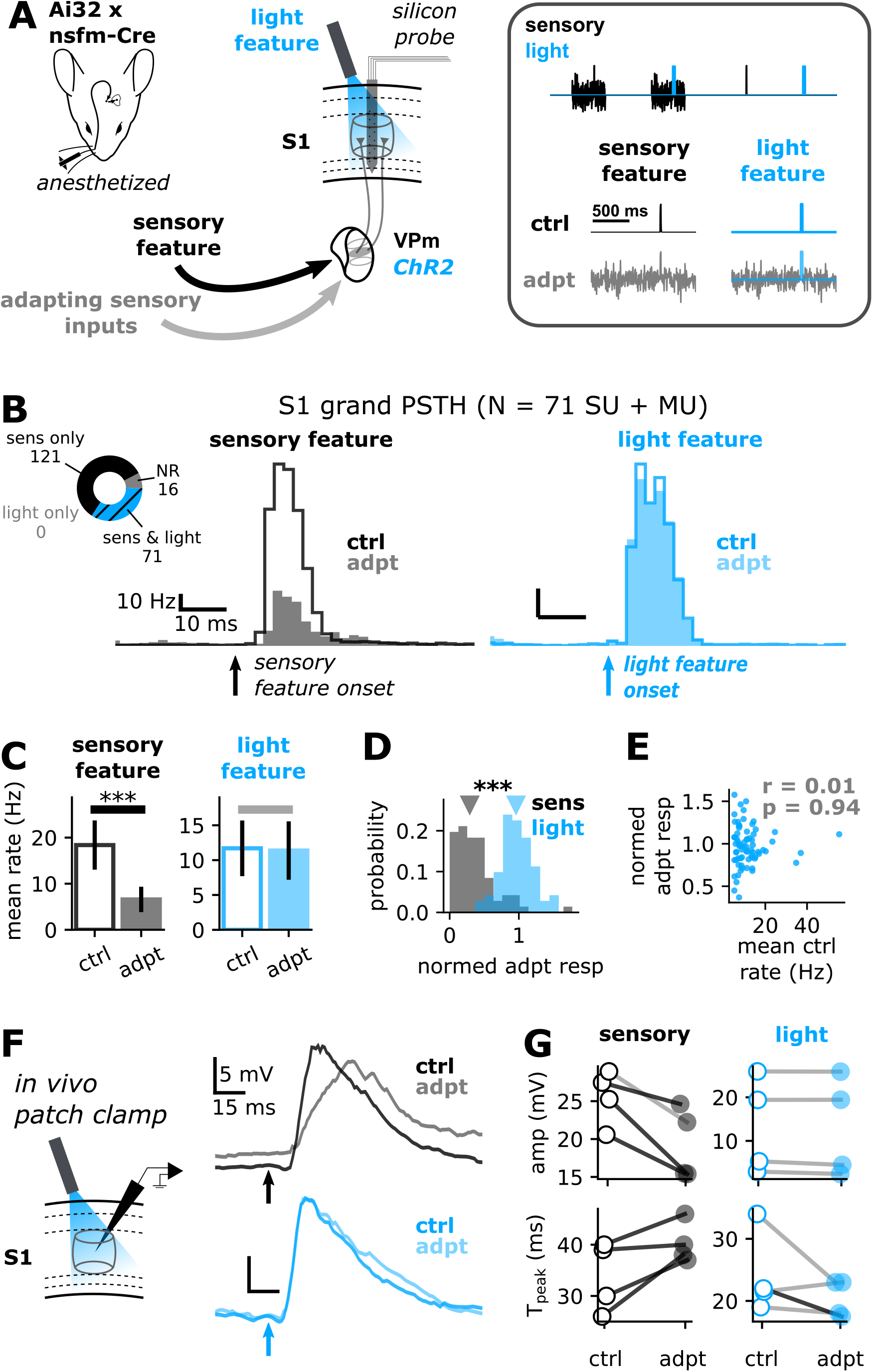
S1 responses to direct optogenetic stimulation of TC terminals are not adapted by sensory white noise. **A.** Experimental setup for extracellular recordings. S1 spiking activity was recorded in mice lightly-anesthetized with isoflurane (see Methods). These transgenic mice expressed Channelrhodopsin in VPm cell bodies, axons, and thalamocortical (TC) axon terminals. An optic fiber positioned above the cortical surface was used to deliver punctate optogenetic stimulation to TC terminals on “light feature” stimulation trials (see Methods). **B.** Grand PSTHs (using all sensory-responsive single- and multi-units, see Methods) for sensory feature (black, left) and light feature (blue, right) trials, for control (empty histogram) and adapted (filled histogram) conditions. Inset: number of recorded units that were significantly responsive to the sensory feature only (“sens only”), the light feature only, both the sensory and light features (“sens & light”), and not responsive to either (“NR”). **C.** Across-unit mean (+/− SEM) punctate-stimulus-evoked firing rates vs stimulus condition for all responsive single- and multi-units, for sensory feature (black, left) and light feature (blue, right) stimuli. (***: p < 0.001, Wilcoxon singed-rank test). Sensory feature mean +/− SEM control: 18.38 +/− 1.27 Hz, adapted: 6.59 +/− 0.65 Hz, W = 39, p = 1.39 × 10^−12^ N = 71 units, Wilcoxon signed-rank test. Light feature mean +/− SEM control: 11.7 +/− 0.95 Hz, adapted: 11.38 +/− 0.99 Hz, W = 987, p = 0.19. **D.** Distributions of normalized adapted responses for all valid units (see Methods). Triangles at top denote population median values (***: p < 0.001, Wilcoxon signed-rank test). Population median normalized adapted response = −0.29 light feature trials, 0.96 sensory feature trials, W = 199, p = 3.12 × 10^−11^, Wilcoxon signed-rank test. **E.** Normalized adapted responses to the light feature vs. across-trial mean light-evoked rate in the control condition for all 71 units that responded to both sensory and light features (with r- and p-values from Pearson correlation test). **F.** Left: experimental setup for in vivo S1 patch clamp recordings in lightly-anesthetized transgenic mice. Right: Across-trial median membrane potential responses to sensory features (black traces) and light features (blue traces), for one example S1 neuron. **G.** Properties of subthreshold responses to sensory features (left) and light features (right): subthreshold response amplitude (top), and time from stimulus onset to peak subthreshold response (T_peak_, center) for each of the four recorded cells. Dark lines connecting pairs of data points indicate significant difference across stimulus conditions (p < 0.05, Wilcoxon signed-rank test), and light lines indicate non-significance.

**Figure 8.**
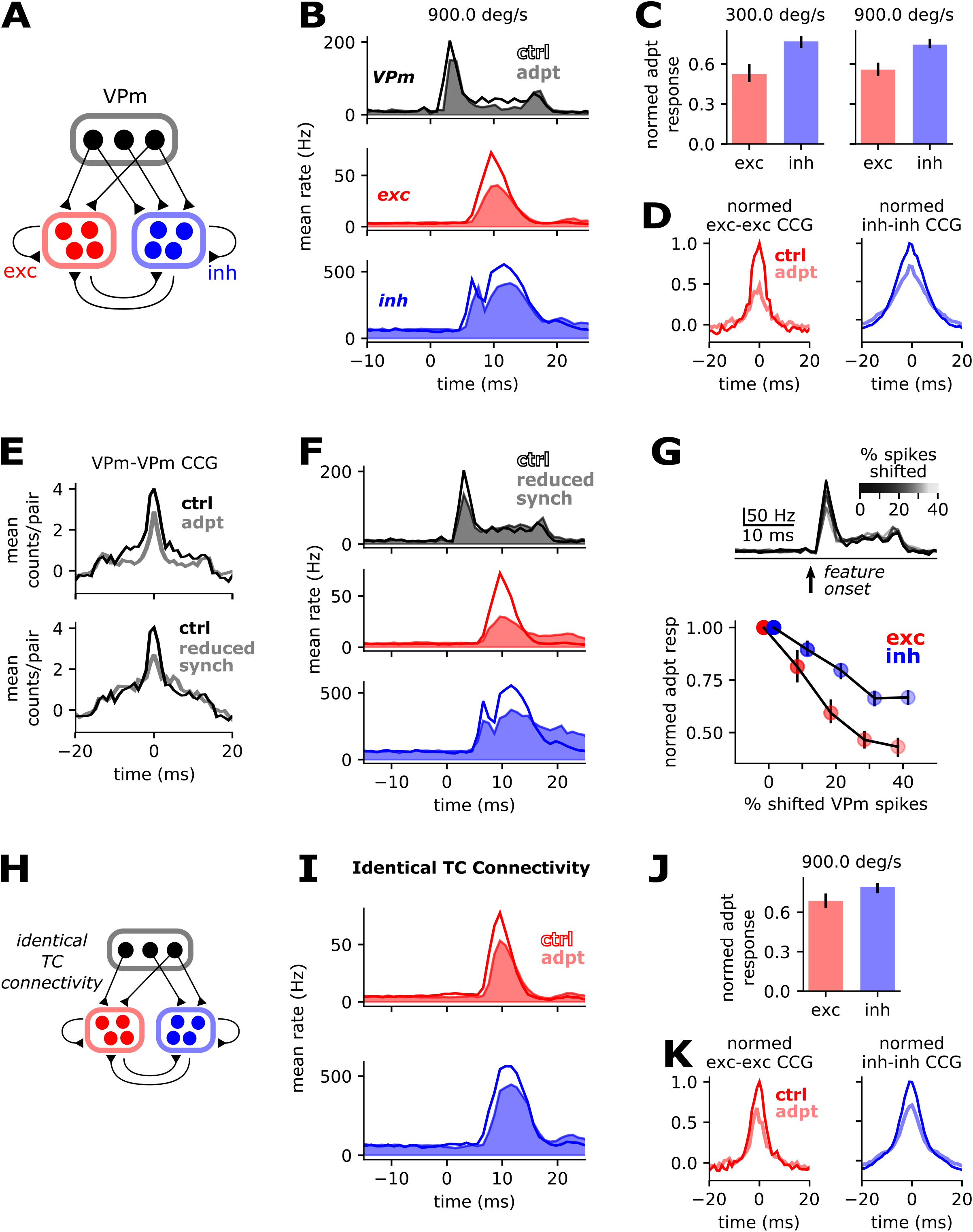
A thalamocortical network model identifies synchronous VPm spikes and feedforward inhibition as key mechanisms underlying response adaptation. **A.** Model schematic (see Methods). **B.** Grand mean PSTHs for VPm spike times used to stimulate the network (top) and network excitatory and inhibitory neurons, for the control (empty PSTHs) and adapted (filled PSTHs) conditions. **C.** Normalized adapted responses for both simulated stimulus velocities (see Methods). For each condition, the response is defined to be the peak of the grand PSTH, and the normalized adapted response is the adapted value divided by the control value. Error bars indicate 95% confidence intervals from re-sampling neurons with replacement. **D.** Grand exc-exc (left) and inh-inh (right) CCGs for 200 randomly-selected pairs of network neurons, for the control (dark line) and adapted (lighter line) conditions. CCGs normalized to max value in control condition for visualization purposes. **E.** Top: grand CCGs for VPm inputs to model in the control (dark line) and adapted (lighter line) conditions (corresponding to PSTHs in B, top). Bottom: grand CCGs for VPm inputs in the control (dark line) and “reduced synch” (lighter line) conditions, where the “reduced synch” condition results from manual changes to drawn VPm spike times (see Methods). **F.** Same as in B, but for control and “reduced synch” simulations. **G.** Top: grand PSTHs for various choices of percent shifted VPm spikes. Bottom: normalized adapted excitatory and inhibitory responses vs. percent of shifted VPm spikes. **H.** Model schematic for “identical TC connectivity” network (see Results and Methods). **I.** Grand excitatory (top) and inhibitory (bottom) PSTHs for “identical TC connectivity” network. **J, K.** Same as in (H, I), but for “identical TC connectivity” network.

### S1 exhibits profound and differential sensory adaptation during wakefulness

We first characterized the effects of the background white noise adapting stimulus on baseline spiking activity in S1. We segregated well-isolated, sensory-responsive cortical units into regular-spiking (RS, putative excitatory) and fast-spiking (FS, putative inhibitory) neurons (Fig. 1B, see Methods). The sensory white noise evoked a sharp increase in mean RS and FS firing, such that mean rates were significantly elevated during the “early” white noise response window relative to spontaneous activity (Fig. 1C, D). Firing rates rapidly adapted, but remained significantly elevated above spontaneous levels in the “late” white noise response window (Fig. 1D). The effect was generally more pronounced for FS cells (Fig. 1C, D). Importantly, this elevation in baseline firing rates did not reflect white-noise-evoked whisking; not only were there distinct features within each PSTH – consistent with stimulus-locked firing and therefore a largely stationary whisker (Fig. 1C) – but we also confirmed using whisker videography in a subset of recording sessions that whisker (Fig. 1D, bottom) and nose (not shown) motion did not systematically change when the whisker white noise was applied (see Methods). As such, it is likely that the induced elevation in cortical firing rates was largely due to afferent sensory drive.

Next, we investigated the adaptive effects of the background stimulus on responses to sensory features during wakefulness. We delivered a 300 deg/s feature to a single whisker, either in isolation (the “control” condition), or embedded in background stimulation (the “adapted” condition, Fig. 1A, see Methods), with the feature delivered 1 s after background stimulus onset. To avoid distortion of the feature waveform, the background stimulus was dampened with an inverted Gaussian waveform in the neighborhood of the feature (Waiblinger et al., 2015; Whitmire et al., 2016; see Methods). This also allowed pre-feature RS and FS firing rates to return to near-baseline values at feature onset (Fig. 1E). We delivered both moderate (300 deg/s) and relatively strong (900 deg/s) features in a subset of experiments. In the control condition, features evoked robust, short-latency spiking responses (Fig. 1E), consistent with previous work in the anesthetized (Bruno and Simons, 2002; Pinto et al., 2003; Khatri et al., 2004; Wang et al., 2010) and awake (Musall et al., 2014) rodent. We next asked whether the background stimulus appreciably adapted feature responses, or whether the relatively high baseline firing rates during wakefulness (Fig. 1D) resulted in a “pre-adapted” circuit (Castro-Alamancos, 2004). We found that S1 feature responses were in fact substantially muted when features were embedded in the background stimulus (Fig. 1E). To characterize RS and FS feature responses and the effects of adaptation, we calculated across-trial mean evoked rates using a 50 ms window following feature onset. For both cell types and feature velocities, the peak (Fig. 1E) and mean (Fig. 1F) evoked rates were reduced in the adapted condition. Interestingly, adaptation appeared to be more profound for RS cells, in terms of proportional changes in sensory responses (Fig. 1E, F). To further quantify the effects of adaptation on a cell-by-cell basis, and to capture cell-type-specific adaptation, we calculated the normalized adapted response for each cell (i.e., the across-trial mean adapted response rate divided by the mean control rate, see Methods). For both feature velocities and cell types, population median normalized adapted responses were less than one (capturing the general adaptive reduction in evoked rate), and RS cells were indeed significantly more adapted than FS cells (Fig. 1G). This differential effect was not specific to any cortical depth (Fig. 1H).

In summary, we observed profound rapid sensory adaptation in S1 of the awake mouse, and cortical putative excitatory neurons were more adapted than putative inhibitory neurons.

### Adaptation reduces the theoretical detectability of punctate sensory stimuli

The adaptation of S1 feature responses should diminish the ability to distinguish the sensory feature from the sensory background. To quantify the potential impact of adaptation on feature detectability, we applied a signal detection theory framework to the spiking activity of putative excitatory neurons, which are likely to play a role in downstream signaling and thus are relevant for percepts. Qualitatively, theoretical feature detectability is inversely related to the degree of overlap between the baseline and feature-evoked firing rate distributions. For each S1 RS unit, we generated spike rate distributions using 50 ms windows of baseline and feature-evoked spiking, and drew samples from “population” gamma distributions (Fig. 2A) parametrized by the empirical mean and standard deviation ((Wang et al., 2010) see Methods). Consistent with the adaptive decrease in single-neuron response rates, adaptation tended to move the feature-evoked distribution toward the baseline distribution, increasing the degree of overlap (Fig. 2A, right). We quantified the overlap by calculating the area under the receiver operator characteristic curve (Wang et al., 2010; Whitmire et al., 2016) (AUROC, Fig. 2B), which has a value of 1.0 for non-overlapping distributions, and 0.5 for complete overlap. Adaptation significantly reduced the across-unit mean AUROC for both feature velocities (Fig. 2C, bottom). Notably, repeating this analysis for a “hybrid” scenario using the adapted feature-evoked distributions and control baseline distributions did not significantly alter AUROC values relative to the true adapted condition (Fig. 2C, left), demonstrating that the decrease in detectability reflected feature response adaptation, rather than an elevation of baseline rates in response to the adapting stimulus.

The above analysis considers spiking in 50 ms pre- and post-feature windows, and the results are consistent with the adaptation of mean feature-evoked RS rates. But it is also apparent from visual inspection of the RS grand PSTHs that the time-course of S1 spiking varied with feature velocity and sensory adaptation. To resolve the dependence of feature detectability on time, we varied the width of the post-feature window from 5 ms to 50 ms in 5 ms increments (see Methods). Consistent with the RS grand PSTHs, we found that in the control condition, AUROC values increased drastically in the first 15 ms of the feature response, and reached peak values within approximately 25 ms (Fig. 2D). The separation between control and adapted AUROC values largely occurred during this early phase of the response (Fig. 2D). In other words, the adaptive loss of theoretical detectability largely reflected a decrease in short-latency feature-evoked spiking.

Together, these results quantify the degree to which sensory adaptation reduces the theoretical detectability of sensory features during wakefulness, and demonstrate that this largely reflects adaptation of the early feature response. This is qualitatively consistent with previous reports showing significant adaptation of theoretical feature detectability in the anesthetized rat (Wang et al., 2010; Ollerenshaw et al., 2014; Zheng et al., 2015), and perceptual threshold in awake rats (Ollerenshaw et al., 2014; Waiblinger et al., 2015).

### Adaptation strongly influences the timing of feature-evoked spiking in S1

As highlighted above, adaptation altered S1 sensory response timing, a key component of cortical sensory representations that strongly influences how and what signals are transmitted to downstream brain structures. Specifically, in addition to the dramatic reduction in early feature-evoked spiking (particularly among RS cells), visual inspection of the grand PSTHs suggested adaptation increased the latency of feature responses. These phenomena may be relevant for interpreting the mechanistic basis of S1 adaptation, and the likely perceptual implications; both are consistent with weaker feedforward excitatory drive to S1, and could influence signaling to downstream targets. We therefore sought to quantify these adaptive effects, and began by calculating the response latency (T_onset_) for each S1 unit (see Methods). We found that for both RS and FS cells, and for both stimulus strengths, adaptation increased T_onset_ values (Fig. 3A). To further quantify and compare adaptive changes in T_onset_ across cell types, we calculated the T_onset_ adaptation index (AI) for each cell (see Methods) and compared population medians for RS and FS cells. For both stimulus strengths, the adaptive increase in response onset times was significantly greater for RS cells (300 deg/s RS median T_onset_ AI = 0.21, FS median T_onset_ AI = 0.02, p = 1.21 × 10^−4^, Kruskal-Wallis test; 900 deg/s RS median T_onset_ AI = 0.13, FS median T_onset_ AI = 0.00, p = 4.93 × 10^−4^).

Next, we reasoned that the observed changes in S1 PSTH shapes might reflect a decrease in synchronous spiking among cortical neurons. In general, the degree of synchronous firing among a population of neurons is likely related to the effect on synaptic targets; stronger inhibitory synchrony will tend to silence postsynaptic neurons, and stronger excitatory synchrony will be more efficacious for driving postsynaptic neurons – possibly downstream from S1. Indeed, it has been shown that coordinated population firing is a better predictor than mean rates of stimulus identity (Jadhav et al., 2009; Safaai et al., 2013; Zuo et al., 2015) and behavioral stimulus discrimination performance (Safaai et al., 2013; Zuo et al., 2015) when rodents whisk across textured surfaces.

To calculate the prevalence of synchronous spikes in feature responses, we populated the grand cross-correlogram (CCG) for RS-RS and FS-FS pairs for each stimulus condition using spike trains from all pairs of simultaneously-recorded responsive cells, and defined synchronous spikes to be those within a +/− 7.5 ms window around zero lag (Fig. 3B, see Methods, Wang et al. 2010). We found that adaptation drastically reduced the amplitude and sharpness of the RS-RS CCGs (Fig. 3C, left), while more modestly reducing the grand CCG amplitude for FS-FS pairs (Fig. 3C, right). This represented a significant decrease in synchronous spike counts for both pair types and stimulus velocities (Fig. 3D). As reflected in the grand CCGs, the decrease in FS-FS synchrony was significant, but proportionally smaller than that of RS-RS synchrony. We next asked whether the adaptive decrease in synchronous spikes simply reflected the decrease in mean evoked rate, or possibly also other de-synchronizing mechanisms. For each feature velocity and cell type, we simulated a “hybrid” condition in which we sub-sampled spikes from the control condition, matching the rate of the adapted condition (see Methods). We found that synchronous spike counts were significantly lower in the adapted than in the hybrid condition (Fig. 3D). Thus, though the relationship between mean rate and synchronous spike count is nonlinear, the adaptive decrease in synchronous spiking was greater than that predicted by the loss of feature-evoked spikes in individual cortical neurons alone. This non-trivial decrease in synchronous firing suggests the feedforward drive to S1 is not only less efficacious for driving spikes in the adapted condition, but may also be less synchronous across thalamorecipient neurons.

Thus, rapid sensory adaptation in S1 during wakefulness not only reduced mean evoked spike rates and theoretical feature detectability, but also disrupted timing through increased response latencies and reduced synchrony. In both aspects of timing, as with mean evoked rates, the adaptive effects on RS cells were more dramatic. The loss of synchronous excitatory firing has implications for the driving of targets downstream of S1, and ultimately for perception and behavior. The more modest decrease in synchronous inhibitory firing implies that inhibitory neurons were still relatively strongly driven in the adapted condition. Further, the synchronous inhibitory spiking that survives adaptation should provide robust feedforward inhibition to S1 excitatory neurons, which may in part explain the more profound adaptation of RS cells.

### S1 sensory responses are also adapted under anesthesia, but RS cells are not differentially adapted

Having established the extent and characteristics of rapid sensory adaptation in S1 of the awake mouse, we next turned our attention to the underlying mechanisms, which may reflect a combination of feedforward, recurrent, and top-down sources. Our first step in doing so was to compare these results to those from the anesthetized mouse, for two reasons. First, while the background stimulus elevated S1 firing rates at least in part via feedforward sensory inputs (as evidenced by the degree of stimulus-locked firing), it is still possible that this adapting stimulus also evoked top-down modulation, e.g., through systematic changes in ongoing S1 state via arousal-related neuromodulation (Reimer et al., 2014, 2016; Mcginley et al., 2015; McGinley et al., 2015). In the anesthetized mouse, ongoing state changes caused by endogenous processes should be independent of the sensory white noise, and as such, white-noise-induced changes in mean feature representations should not reflect top-down mechanisms. Second, isoflurane anesthesia tends to silence the secondary posteromedial nucleus of thalamus (Suzuki and Larkum, 2020), lowers baseline cortical firing rates (Greenberg et al., 2008; Vizuete et al., 2012; Aasebø et al., 2017), and generally weakens cortical inhibition (Haider et al., 2013) and other intracortical interactions (Suzuki and Larkum, 2020). We therefore repeated our experiments in a different set of mice lightly-anesthetized with isoflurane, using the anesthesia to unmask the feedforward inputs from VPm to S1.

As expected, baseline firing rates under anesthesia were quite low compared to those recorded during wakefulness (Fig. 4B, C). Despite these differences in baseline activity, the anesthetized experiments recapitulated several key aspects of the awake data. First, the background stimulus evoked a sharp increase in RS and FS firing in the “early” white noise response window, with rapid adaptation toward “late” response rates that were significantly elevated above spontaneous values (Fig. 4B, C), and with stimulus-locked firing evident among FS neurons (Fig. 4B). Further, adaptation clearly decreased the peak (Fig. 4D) and mean evoked firing rates of responses to 300 deg/s features (Fig. 4E), and qualitatively shifted PSTHs to higher response latencies (Fig. 4D). Notably, however, excitatory neurons were not differentially adapted under anesthesia (Fig. 4F). In light of this, it is possible that the stronger cortical inhibition typical of wakefulness (Haider et al., 2013) contributed to adaptation of RS cells in our awake recordings via robust feedforward inhibition.

In summary, the elevation of (stimulus-locked) firing with presentation of background sensory stimulation, and net adaptation of S1 feature responses was robust to anesthesia, suggesting that feedforward mechanisms are sufficient to explain these phenomena. In contrast, the differential adaptation of S1 excitatory neurons was abolished by anesthesia, suggesting the set of mechanisms underlying this phenomenon possibly including strong feedforward inhibition – were not all active in the anesthetized state.

### Adaptation primarily influences feature response timing in VPm thalamus

Having established that several aspects of S1 sensory adaptation may reflect feedforward mechanisms, we next sought to identify those mechanisms. Previous *in vitro* and anesthetized work has demonstrated adaptation of evoked rate (Hartings et al., 2003; Khatri et al., 2004; Gabernet et al., 2005; Ganmor et al., 2010; Wang et al., 2010; Ollerenshaw et al., 2014; Whitmire et al., 2016; Liu et al., 2017), single-unit bursting (Whitmire et al., 2016) and population synchrony (Wang et al., 2010; Ollerenshaw et al., 2014) in VPm, suggesting S1 adaptation is inherited to some degree directly from VPm, yet this has never been tested during wakefulness. We therefore next repeated our experiments while recording spiking activity in VPm of the awake mouse (Fig. 5A, see Methods), and asked how adaptation altered the rate and timing of thalamic sensory responses.

We first parsed VPm spikes into putative burst and tonic spikes, and asked how presentation of sensory white noise affected baseline tonic and burst firing. We defined bursts according to classic criteria (Lu et al., 1992; Reinagel et al., 1999; Swadlow and Gusev, 2001; Lesica et al., 2006; Whitmire et al., 2016): two or more sequential spikes from a single unit preceded by at least 100 ms of quiescence, and with at most 4 ms between spikes (Fig. 5A, bottom). These criteria are consistent with the timing of burst spikes resulting from de-inactivation of T-type calcium channels after prolonged hyperpolarization (McCormick, 1992; Llinás and Steriade, 2006), though we did not confirm this directly via intracellular recordings. As in S1, we found that the background stimulus evoked stimulus-locked firing in VPm (Fig. 5B). Tonic firing rates increased sharply at white noise onset, and rapidly adapted to “late” white noise response rates that were significantly elevated above spontaneous levels (Fig. 5B, C). Burst rates were very low during spontaneous activity (Fig. 5B, C), increased sharply at white noise onset, before decreasing below spontaneous levels in the “late” response window (Fig. 5B, C) – consistent with inactivation of T-type calcium channels. We next characterized VPm responses to sensory features, and the effects of the background stimulus on these responses. We found that background stimulation reduced feature-evoked tonic spiking (Fig. 5D, E), but this adaptive effect was modest in comparison to the adaptation of S1 RS cells. Adaptation had a more striking effect, however, on burst firing in individual neurons. Burst firing has been shown to provide potent synaptic drive to S1 (Sherman, 2001; Swadlow and Gusev, 2001; Swadlow, 2002), and may therefore be critical for shaping cortical sensory responses. Yet few studies have explored single-unit VPm sensory responses during wakefulness, when VPm is likely to be on average relatively depolarized, and T-type calcium channels inactivated. We did, in fact, observe feature-evoked bursting, consistent with previous recordings in awake rats (Whitmire et al., 2016), though burst spikes constituted a minority of total feature-evoked spikes in the control condition (Fig. 5E). Adaptation profoundly reduced bursting, with feature-evoked burst spikes nearly abolished in the adapted condition (Fig. 5D, E). Finally, we asked how adaptation altered synchronous thalamic firing, shown previously to be crucial for driving downstream cortical targets (Bruno and Sakmann, 2006; Wang et al., 2010; Bruno, 2011). We thus generated grand CCGs and counted synchronous spikes for each stimulus condition, as done previously for S1 cells. We found that adaptation significantly and substantially decreased synchronous VPm spike counts for both feature velocities (Fig. 5F, G). Interestingly, adapted synchronous spike counts were not significantly different from those of the hypothetical “hybrid condition” (Fig. 5G). Thus, unlike S1, VPm was apparently not subject to mechanisms that actively de-synchronized firing across thalamic neurons beyond the degree predicted by loss of rate alone. As such, the loss of synchrony in this regime of adaptation may be attributable to weakened synaptic drive from trigeminal neurons (Castro-Alamancos, 2002; Timofeeva and Lavalle, 2003; Ganmor et al., 2010). This is in contrast to previous studies in anesthetized (Wang et al., 2010) and awake (Ollerenshaw et al., 2014) rats, which demonstrated that sequences of high-velocity stimulus features resulted in a gradual loss of synchrony greater than that predicted by adaptation of mean rate alone. This discrepancy may owe to the relatively weak adapting stimulus we employ here; stronger adapting stimuli may engage a broader range of mechanisms that serve to desynchronize VPm firing, possibly including more profound adaptation of trigeminal neurons, and weakened synaptic interactions between VPm and TRN.

In summary, rapid sensory adaptation modestly reduced feature-evoked tonic spiking in VPm, and substantially reduced bursting and synchronous spiking. Given the sensitivity of S1 to bursting and synchronous firing in VPm (Usrey et al., 2000; Sherman, 2001; Swadlow and Gusev, 2001; Swadlow, 2002; Bruno and Sakmann, 2006; Wang et al., 2010; Ollerenshaw et al., 2014), this qualitatively predicts that cortical responses will be smaller in amplitude and less synchronous across neurons, as observed in experiment. Further, the more substantial loss of synchronous firing in S1 than in VPm demonstrates an apparent enhancement of adaptation-induced desynchronization from thalamus to cortex.

### Thalamocortical and intracortical synaptic depression contribute little to S1 sensory adaptation

While the thalamic adaptation we observed likely played a key role in S1 adaptation, what was the additional contribution from thalamocortical and/or intracortical synaptic depression? Previous anesthetized work has demonstrated greater adaptation of mean evoked rates in S1 than in VPm (Chung et al., 2002; Khatri et al., 2004; Gabernet et al., 2005), and this has been interpreted to reflect thalamocortical synaptic depression. However, this may also reflect the sensitivity of cortex to bursting and synchronous thalamic firing, which we found to be profoundly adapted by persistent sensory stimulation. We sought to disentangle the relative contributions of thalamic response adaptation and thalamocortical and/or intracortical synaptic depression by performing two complementary optogenetic experiments: one to isolate the contribution from thalamocortical synaptic depression, and one to isolate the contribution from intracortical mechanisms. In these experiments, we achieved targeted manipulation of either VPm neurons or thalamocortical synapses – without directly manipulating whisker-responsive POm or TRN neurons – by using a transgenic mouse (Ai32xNR133, see Methods) expressing Channelrhodopsin in VPm/VPl cell bodies, axons, and thalamocortical axon terminals.

In the first experiment, we sought to isolate the contribution of thalamocortical synaptic depression to S1 response adaptation. We did this by presenting a background optogenetic stimulus that bypassed the early sensory pathway upstream of VPm, and elevated baseline VPm rates without elevating S1 rates. We inserted an optoelectrode into VPm and a linear silicon probe into the topographically-aligned column of S1 in the awake mouse (Fig. 6A, see Methods), and we randomly interleaved “white noise” and “LED” trials. On “LED” trials, we elevated baseline VPm rates by substituting the adapting sensory stimulus with a step input of blue light to thalamus (Fig. 6B, right). We titrated the light power such that mean baseline thalamic rates were comparable to “white noise” trials (Fig. 6B, 6C, top). Importantly, while the LED significantly increased VPm firing rates (Fig. 6B, C top, spont vs. LED), it did not evoke even a transient increase in S1 firing rates (Fig. 6B, C, bottom), likely because thalamic spiking was not sufficiently synchronous to effectively drive cortical targets (Bruno and Sakmann, 2006). As such, this manipulation did not engage activity-dependent intracortical adaptation mechanisms.

We next inspected the effects of our sensory and optogenetic manipulations on VPm and S1 sensory feature responses. We were interested in the presence or absence of gross adaptive effects, and so we grouped together tonic and burst spikes in VPm, and RS and FS cells in S1 for this analysis. Because sensory adaptation was qualitatively similar for 300 deg/s and 900 deg/s features, we presented only the 300 deg/s sensory feature in these experiments. On LED trials, we maintained a constant light level during presentation of the sensory feature, to avoid transient VPm responses to reduction in light power. As shown above, background sensory stimulation adapted feature responses in VPm and S1 (Fig. 6D). We next inspected the effects of optogenetically-elevated baseline VPm rates on feature responses. If the “artificial” elevation of baseline VPm rate adapted TC synapses prior to delivery of the sensory feature, we would anticipate adapted S1 feature responses on LED trials, despite the non-adapted VPm feature response. On the contrary, we observed no significant differences in S1 feature response rates between the control and LED conditions (Fig. 6D, bottom). Importantly, LED presentation did not significantly enhance synchronous spike counts in the VPm feature response relative to control trials (Fig. 6E), thus ruling out the possible confound of enhanced feature-evoked thalamic synchrony masking the effects of TC synaptic depression. As such, the results of this first optogenetic experiment suggest that S1 response adaptation did not largely reflect TC synaptic depression, but rather a combination of VPm response adaptation and intracortical mechanisms.

We next performed a second, complementary optogenetic experiment to determine the relative contribution from intracortical mechanisms engaged by the elevation of pre-feature cortical firing rates (e.g., intracortical synaptic depression, a build-up of intracortical inhibition, etc.). We positioned an LED-coupled optic fiber above the cortical surface and recorded extracellular spiking activity with a silicon probe array in S1 (Fig. 7A). We performed these experiments in lightly-anesthetized mice – in which we also observed profound S1 adaptation (Fig. 4) – which allowed additional recording time to titrate light levels (see Methods). On each trial, we presented either a 300 deg/s sensory feature to the principal whisker (as described above), or a “light feature”, which was a brief step input of light to stimulate TC terminals in the principal column, with sensory and light features randomly interleaved. In both cases, we presented the features either in isolation (control trials) or embedded in a background sensory stimulus (adapted trials). We recorded from a pool of 208 RS, FS, and multi-units across these experiments. Of these, 121 responded significantly to the sensory but not the light feature, 0 responded only to the light feature, 71 responded to both, and 16 did not respond to either (Fig. 7B, inset). For the following analyses, we consider only the units that responded significantly to both feature types (see Methods).

Consistent with the above results for awake and anesthetized mice, the S1 sensory feature response grand PSTH for pooled RS, FS, and multi-units exhibited profound sensory adaptation (Fig. 7B, left). If this largely reflected intracortical mechanisms engaged prior to feature delivery, we would expect that light feature responses to be similarly adapted by the background sensory stimulus, as the light feature precluded VPm response adaptation. Instead, there was comparatively little evidence of adaptation in the S1 grand PSTHs for light feature responses (Fig. 7B, right). This result also bore out in mean evoked rates: mean rates for sensory feature responses were profoundly adapted (Fig. 7C, left), while light feature responses were only slightly adapted (Fig. 7C, right), and the population median normalized adapted response was near 1 for light feature responses, but significantly smaller for sensory feature responses (Fig. 7D).

Importantly, this did not appear to simply reflect an overwhelmingly strong LED stimulus, or light-evoked TC synaptic activity that was unnaturally synchronous across synapses; not only were evoked rates generally lower for the light feature than for the sensory feature across all neurons (Fig. 7B, C), but in exploring a variety of LED stimulus amplitudes and durations across experiments, we found that both relatively large- and small-amplitude light-evoked responses were at most only modestly adapted, and that the degree to which sensory white noise adapted light-evoked responses was unrelated to the amplitude of the control response for a given unit (Fig. 7E). Further, given that we observed no units that were responsive to the light but not the sensory feature (Fig. 7B, inset), it is also unlikely that the light feature responses were driven primarily by TC synapses and/or cortical neurons that were not responsive to sensory stimulation (and therefore could not be adapted by sensory white noise).

We confirmed these observations of spiking activity by obtaining in vivo patch clamp recordings from neurons in S1 of the lightly-anesthetized mouse (Fig. 7F, left, see Methods). We recorded from four neurons that responded to both sensory and light features. While sensory- and light-evoked amplitudes varied across neurons (Fig. 7G, top), light feature responses were comparatively impervious to sensory adaptation. Specifically, for sensory feature responses, the across-trial median amplitude significantly decreased for three cells, and the time to response peak significantly increased for all cells, (Fig. 7G, left, see Methods), consistent with the extracellular recordings (Fig. 7B, C). In contrast, light feature response amplitudes were unchanged and the time to peak response did not increase for any of these cells (Fig. 7G, right).

These two complementary optogenetic experiments together suggest that intracortical mechanisms engaged during pre-feature activity – including TC synaptic depression – contributed little to S1 response adaptation, which in this regime of adaptation appear to primarily reflect the profound loss of synchronous feature-evoked VPm spiking.

### A model network identifies synchronous VPm spikes and robust feedforward inhibition as key mechanisms underlying S1 response adaptation

We observed that S1 feature response rates were more profoundly adapted than VPm feature response rates, and we have proposed that this reflected the sensitivity of S1 to the adaptive loss of synchronous VPm spikes. Further, we hypothesized that feedforward cortical inhibition that was relatively robust to sensory adaptation helped to dampen feature-evoked S1 excitatory firing in the adapted condition. In other words, inhibition increased the sensitivity of cortical excitatory neurons to adaptation of synchronous VPm spiking. To test the reasonableness of these assertions, and to identify the mechanistic basis for robust feedforward inhibition, we implemented a model thalamocortical network, and assessed its ability to reproduce profound, cell-type-specific adaptation.

We modeled a single S1 barrel as a clustered network of 800 excitatory and 100 inhibitory leaky integrate-and-fire neurons, subject to excitatory inputs from a model “VPm barreloid” (Fig. 8A). The barreloid was modeled as 40 independent trains of tonic and burst spikes, with spike times drawn from the empirical VPm PSTHs. We selected cortical network and intrinsic neuronal parameters that mimicked measurements from previous studies, and then adjusted parameters slightly to ensure stable ongoing and evoked network activity (see Methods). Importantly, we implemented differential thalamocortical (TC) connectivity, which we hypothesized might contribute to the robustness of inhibitory firing to VPm adaptation. This differential connectivity consisted of three key components motivated by previous experimental work. First, inhibitory neurons had higher “TC convergence” (or proportion of VPm neurons that synapse onto each cortical neuron) than excitatory neurons (0.75 vs. 0.5, (Bruno and Simons, 2002)). Second, the baseline and evoked firing rate of each VPm neuron was drawn from a skewed distribution, and VPm neurons with the highest rates synapsed exclusively onto inhibitory neurons ((Bruno and Simons, 2002), see Methods). Finally, TC synaptic latencies were on average 1 ms shorter for inhibitory neurons (Cruikshank et al., 2007; Kimura et al., 2010). With this architecture in place, we modeled 50 trials from each stimulus condition of interest.

We fine-tuned the model parameters to give qualitatively realistic peak rates for excitatory neurons when VPm spike times were drawn from the control (unadapted) empirical PSTHs for the 300 deg/s and 900 deg/s sensory features (Fig. 8B). Thus, the excitatory population was appropriately tuned to the rate and synchrony of thalamic firing. Next, we repeated the simulations using VPm spikes drawn from the adapted PSTHs (Fig. 8B, top, filled PSTH), and found that cortical network excitatory neurons were in fact profoundly adapted, despite the only modest reduction in mean VPm rates (Fig. 8B, center). Further, the excitatory population was more strongly adapted than the inhibitory population (Fig. 8B, C). Finally, as observed in experiment, synchronous spike counts were significantly reduced in the adapted condition for excitatory-excitatory (Fig. 8D, left) and inhibitory-inhibitory (Fig. 8D, right) pairs (see Methods), with a more drastic reduction for excitatory neurons. Thus, the mechanisms incorporated in this simple model were sufficient to qualitatively reproduce our key experimental results.

In these simulations, both the mean VPm rate (Fig. 8B, top) and VPm synchronous spiking (Fig. 8E, top) were reduced in the adapted condition. We next assessed the degree to which the loss in synchronous VPm spikes alone could explain cortical adaptation. To do this, we repeated the simulations while “manually” manipulating VPm spike times. Specifically, we first drew VPm spike times from the control PSTHs, and for each spike that occurred within 5 ms of the PSTH peak time, we shifted the spike to a random higher latency (within approximately 20 ms of the peak) with 30% probability (see Methods). This had the effect of maintaining the mean evoked VPm rate, while reducing the number of near-coincident pairs of VPm spikes in the early response (Fig. 8E, bottom, “reduced synch” condition). We found that this change alone – which only modestly affected the resulting VPm grand PSTH (Fig. 8F, top) – was sufficient to profoundly adapt mean excitatory evoked rates (Fig. 8F). Further, excitatory and inhibitory responses decreased monotonically with percentage of shifted VPm spikes, with excitatory cells more sensitive to this manipulation (Fig. 8G). In other words, a loss of synchronous VPm spiking alone was sufficient to reproduce profound and differential cortical response adaptation in this network.

Finally, we asked whether robust feedforward inhibition – mediated in part by differential TC connectivity – contributed to the adaptation of network excitatory neurons. We modified the network slightly by setting identical TC convergence and TC synaptic latency values for excitatory and inhibitory neurons and eliminating rate-dependent TC connectivity (Fig. 8H). Inhibitory and excitatory neurons therefore had identical mean thalamocortical connection properties, though differences in intrinsic neuronal properties and dense excitatory-to-inhibitory connectivity still allowed for higher mean firing rates in the inhibitory population (see Methods). We then slightly reduced the mean TC synaptic weight to yield reasonable excitatory responses in the control condition (see Methods), before inspecting the responses to adapted VPm inputs. For this network, evoked rates (Fig. 8I, top, J) and synchronous spike counts (Fig. 8K, left) for excitatory neurons were only modestly adapted compared to the model with differential TC connectivity, and the degree of adaptation more closely matched that of the inhibitory population (Fig. 8J, K, right). In other words, the excitatory population was less sensitive to VPm adaptation when differential TC connectivity was removed. We used an additional set of models to further assess the relative importance of each component of the differential TC connectivity in the original model. While each component contributed, we found that the degree of excitatory adaptation was most sensitive to differences in TC synaptic latencies (not shown). Importantly, imposing identical TC connectivity did not entirely remove differential adaptation, as the intrinsic inhibitory neuronal properties and dense intracortical connectivity also helped make them comparatively robust to losses in synchronous spiking in both thalamic and cortical neurons excitatory neurons (not shown). This model thus demonstrates the role of robust feedforward inhibition – due in part to differential TC connectivity – in shaping the adaptation of cortical excitatory neurons.

Taken together, these modeling results support our hypotheses that the profound adaptation of cortical RS cells during wakefulness represents a loss of synchronous feature-evoked thalamic spikes, in conjunction with strong feedforward inhibition that is comparatively robust to this decrease in feedforward thalamic drive.

## DISCUSSION

To determine the nature of rapid cortical sensory adaptation during wakefulness, we recorded single-unit activity in S1 of the awake, head-fixed mouse while presenting punctate sensory features either in isolation, or embedded in a persistent background stimulus. To elucidate the mechanistic basis of cortical adaptation, we recorded from putative excitatory and inhibitory cortical neurons and the lemniscal inputs to S1, while employing a battery of additional manipulations across the thalamocortical circuit. This approach, in conjunction with a thalamocortical network model, allowed us to infer the contributions from thalamic adaptation, thalamocortical synaptic depression, and intracortical mechanisms.

Previous in vitro and anesthetized work has clearly demonstrated profound rapid sensory adaptation in sensory cortex (Lesica et al., 2007; Heiss et al., 2008; Ganmor et al., 2010; Wang et al., 2010; Cohen-Kashi Malina et al., 2013; Ollerenshaw et al., 2014; Zheng et al., 2015; Kheradpezhouh et al., 2017), which is thought to represent the net effects on the circuit of elevated firing rates. Yet because baseline cortical firing rates are elevated during wakefulness compared to the anesthetized state (Greenberg et al., 2008; Vizuete et al., 2012; Aasebø et al., 2017), it remains an open question how much room is left for fine-tuning by the sensory environment (Castro-Alamancos, 2004). Here, we demonstrate that cortical sensory responses can indeed be profoundly adapted during wakefulness. The adaptive decrease in theoretical feature detectability and synchronous firing of putative cortical excitatory neurons suggest that downstream targets of S1 will be substantially less driven in the adapted state, which predicts a decrease in perceived feature intensity and a loss of detectability. These observations are consistent with previous behavioral work in rats, which demonstrated changes in perceptual reporting following repetitive whisker stimulation (Ollerenshaw et al., 2014; Waiblinger et al., 2015). In other words, this study supports previous in vitro and anesthetized work suggesting cortical response adaptation could underlie perceptual adaptation.

Our observations are also consistent with elements of two previous studies in awake rats, which demonstrated adaptation of S1 spiking (Musall et al., 2014) and LFP deflections (Castro-Alamancos, 2004) during presentation of feature sequences across a range of animal states. Interestingly, Castro-Alamancos (2004) showed that the extent of adaptation varied across behavioral states, from profound during quiescence to nearly absent in the very early stages of training in an active avoidance task (i.e., likely during states of hyper-arousal). Musall et al. (2014), on the other hand, showed that the degree of cortical response adaptation was largely unchanged by task engagement. This suggests that sensory adaptation is likely relevant across much of the continuum of behavioral states, and that brain state gates adaptation, but future work is needed to more fully address the relationship between behavioral state and adaptation. Active sensation is a particularly important and complex scenario: whisker self-motion may invoke sensory adaptation via bottom-up inputs due to re-afference (Yu et al., 2016) and low-amplitude whisker deflections resulting from surface contacts, but is also associated with top-down modulation of VPm and S1 state (Poulet et al., 2012; Poulet and Crochet, 2019). While our study provides insight into how isolated bottom-up inputs may adapt sensory representations and percepts, it is not clear how whisking-related state modulation will gate this process.

We next sought to identify the mechanistic basis for S1 response adaptation. One body of literature implicates thalamocortical synaptic depression (Castro-Alamancos and Oldford, 2002; Chung et al., 2002; Gabernet et al., 2005; Cruikshank et al., 2010), while our previous work points to adaptation of thalamic spike timing (Wang et al., 2010; Ollerenshaw et al., 2014; Whitmire et al., 2016), but both viewpoints have originated largely from in vitro or anesthetized preparations. Our results suggest cortical response adaptation is largely due to thalamic response adaptation in the regime we explored here. First, adaptation profoundly reduced single-unit bursting and the rate of feature-evoked synchronous spikes in VPm, likely due to a combination of persistent VPm depolarization (Whitmire et al., 2016) and adapted inputs from trigeminal neurons (Castro-Alamancos, 2002; Timofeeva and Lavalle, 2003; Ganmor et al., 2010), which predicts attenuated cortical firing (Swadlow and Gusev, 2001; Bruno and Sakmann, 2006; Wang et al., 2010; Ollerenshaw et al., 2014). Further, optogenetic elevation of baseline VPm rates did not adapt S1 feature responses, and background sensory stimulation had little effect on S1 responses to direct TC terminal stimulation. Finally, our modeling demonstrated that modest reductions in synchronous VPm spiking alone predicted profound cortical adaptation. Taken together, these results therefore demonstrate for the first time in the awake animal the sensitivity of cortex to thalamic spike timing in the context of sensory adaptation, and suggest that synaptic depression contributes little to the observed S1 feature response attenuation.

This apparent lack of TC synaptic depression appears to contradict the results of previous anesthetized and in vitro studies (Castro-Alamancos and Oldford, 2002; Chung et al., 2002; Gabernet et al., 2005; Cruikshank et al., 2010). We believe that this reflects a difference in the strength of adapting stimuli. Specifically, these previous studies used adapting sequences of high-velocity, punctate sensory stimuli and/or electrical stimulation. Here, we employed a relatively low-amplitude white noise adapting stimulus, to emulate small-amplitude whisker micromotions that occur during whisking across textured surfaces (Jadhav and Feldman, 2010). In terms of total power (Zheng et al., 2015), this white noise stimulus is many times weaker than the adapting stimuli used in many previous studies (not shown). This likely adapts the TC circuit to a lesser degree than punctate stimulus trains, which may explain the apparent contradiction with previous work. This may also explain why we did not observe more pronounced adaptation of FS units, which has been shown to reflect stronger adaptation of TC synapses onto inhibitory neurons (Gabernet et al., 2005). It is possible that stronger adapting stimuli will generally engage a broader range of adaptive mechanisms than we observed here, with thalamocortical/intracortical synaptic depression playing more prominent roles in regimes of more profound adaptation. It is also unclear whether sensory white noise presented during wakefulness can alter aspects of cortical sensory coding not explored here (such as input-output relationships (Maravall et al., 2007) and response tuning curves (Wang et al., 2010)), and whether adaptation of thalamic synchrony is sufficient to explain such changes. Future work probing a range of adapting stimulus strengths and effects can investigate such questions.

While thalamic feature response adaptation appeared necessary for S1 adaptation, it did not explain the differential adaptation of RS and FS cells during wakefulness. In other words, RS and FS cells provided different read-outs of feature-evoked thalamic spiking. In contrast, RS feature responses were no more adapted than FS rates under isoflurane anesthesia, which has been shown to disproportionately weaken cortical inhibition (Haider et al., 2013; Taub et al., 2013). This suggested to us that feedforward inhibition contributed to the adaptation of RS cells during wakefulness. We explored this possibility with a network model, in which we implemented cell-type-specific TC connectivity motivated by previous experimental work (Bruno and Simons, 2002). We found that S1 response adaptation did largely reflect a loss of synchronous VPm spikes, but that the profound and differential adaptation of excitatory neurons also required several additional mechanisms, including cell-type-specific TC connectivity. Taken together, these experimental and modeling results suggest a thalamocortical circuit basis for the observed S1 adaptation, involving a profound loss of synchronous feedforward excitation, and a comparatively modest decrease in dampening feedforward inhibition.

This adaptive shift in the feedforward E/I balance toward inhibition has implications for cortical function and perception beyond attenuation of response amplitudes and perceived stimulus intensities. For example, previous experimental work has demonstrated that the relative strength and/or timing of cortical excitation and inhibition contributes to the direction-selectivity (Wilent and Contreras, 2005) and receptive field properties (Kyriazi and Simons, 1993; Bruno and Simons, 2002; Ramirez et al., 2014) of excitatory neurons, maintains relatively low excitatory firing rates during bouts of whisking (Yu et al., 2016; Gutnisky et al., 2017), shapes the “window of integration” during which excitatory neurons integrate excitatory synaptic inputs and depolarize toward threshold (Gabernet et al., 2005; Wilent and Contreras, 2005), and generally serves to “dampen” thalamic-evoked spiking in the excitatory subnetwork (Pinto et al., 2003). Thalamocortical adaptation exists on a continuum (Wang et al., 2010; Zheng et al., 2015), and more moderate levels of adaptation than we imposed here may result in moderately attenuated excitatory firing that is sharpened in space and time by comparatively non-adapted inhibition, resulting in more faithful spatiotemporal cortical representations of complex sensory stimuli. Future experiments exploring regimes of weaker adaptation and using more complex single- and multi-whisker stimulation can explore these possibilities.

In summary, these results demonstrate the effects of rapid sensory adaptation on the early sensory pathway during wakefulness, culminating in profound adaptation of primary sensory cortex, which likely underlies perceptual adaptation on this timescale. Further, they highlight the relative importance of thalamic gating in establishing cortical adaptation, through population timing control of thalamic drive and the differential engagement of the inhibitory cortical sub-population.

## Acknowledgements

We thank Aurelie Pala, Audrey Sederberg, Adam Willats, Christian Waiblinger, and Elaida Dimwamwa for helpful comments on the data analysis and manuscript. We thank Aurelie Pala for assistance with spike-sorting, and Mahmood S. Hoseini for his assistance with the network model. **Funding**. This work was supported by NIH National Institute of Neurological Disorders and Stroke BRAIN Grant R01NS104928 (GBS), NIH National Institute of Mental Health BRAIN Grant U01MH106027 (GBS and CRF), NIH National Institute of Neurological Disorders and Stroke Pre-doctoral NRSA NS098691 (PYB), NIH National Eye Institute R01EY023173 (CRF), and National Science Foundation Graduate Research Fellowships (WMS, MFB).

